# Testing evolutionary explanations for the lifespan benefit of dietary restriction in *Drosophila melanogaster*

**DOI:** 10.1101/2020.06.18.159731

**Authors:** Eevi Savola, Clara Montgomery, Fergal M. Waldron, Katy Monteith, Pedro Vale, Craig Walling

## Abstract

Dietary restriction (DR), limiting calories or specific nutrients, extends lifespan across diverse taxa. This lifespan extension has been explained as diet-mediated changes in the trade-off between lifespan and reproduction, with survival favoured with scarce resources. Another evolutionary hypothesis suggests the selective benefit of the response is the maintenance of reproduction. This hypothesis predicts that lifespan extension is a side effect of benign laboratory conditions, where DR individuals are frailer and unable to deal with additional stressors, and thus lifespan extension should disappear under more stressful conditions. We tested this by rearing outbred female *Drosophila melanogaster* on 10 different protein:carbohydrate diets. Flies were either infected with a bacterial pathogen (*Pseudomonas entomophila*), injured or unstressed. We monitored lifespan, fecundity and ageing measures. DR extended lifespan and reduced reproduction irrespective of injury and infection. These results do not support lifespan extension under DR being a side effect of benign laboratory conditions.

## INTRODUCTION

Nutrition has long been of interest in the field of aging research, particularly due to its potential applications to an ageing human population (reviewed in Bertozzi et al., 2016; Redman & Ravussin, 2011; Speakman & Mitchell, 2011). Dietary restriction (DR), the limitation of a particular nutrient or the overall caloric intake, has been shown to extend lifespan and delay ageing across a range of organisms (reviewed in Mair & Dillin, 2008). Its prevalence and taxonomic diversity suggests the response is evolutionarily conserved and acts via conserved mechanisms (reviewed by Fontana et al., 2010). As such, a large body of research has focused on using the DR paradigm to try to understand the mechanisms underlying variation in ageing and lifespan (e.g. Fontana & Partridge, 2015; Gems & Partridge, 2012; Gibbs & Smith, 2016). However, the evolutionary basis of the response has been much less well investigated (Raubenheimer et al., 2016; Regan et al., 2020; Travers et al., 2020; Zajitschek et al., 2016). This is surprising given that knowledge of the evolutionary basis of the DR response is important to understanding under what conditions it may be applicable in human health. Here we test the two main evolutionary explanations for lifespan extension under DR, which make contrasting predictions about how this response should vary across environments.

The predominant evolutionary explanation, termed the resource reallocation hypothesis (RRH) (Adler & Bonduriansky, 2014; Regan et al., 2020), explains the observed DR response as an adaptive shift in relative investment of resources into survival versus reproduction (Adler & Bonduriansky, 2014; Kirkwood, 1977; Shanley & Kirkwood, 2000). A food shortage signals a sub-optimal environment, where the number and survival probability of any offspring produced is likely to be low (Holliday, 1989; Shanley & Kirkwood, 2000). Under such conditions, an individual could maximise fitness by temporarily delaying reproduction and instead investing resources into survival and somatic maintenance. Once food availability returns, the individual could then maximise fitness by investing resources back into reproduction. By maintaining individuals chronically on low food, aging rates decrease and the individual lives longer (Holliday, 1989; Shanley & Kirkwood, 2000). The RRH requires a trade-off between investing resources into reproduction versus somatic maintenance (Holliday, 1989) and that the response evolved in an environment which fluctuates between low and high food availability (Adler & Bonduriansky, 2014).

In contrast to the predictions of the RRH, some studies suggest that survival and reproduction can be uncoupled under DR (Flatt, 2011). In addition, wild systems have much higher levels of extrinsic mortality than laboratory conditions (for example, from predators or disease), potentially making an individual less likely to live long enough to benefit from delayed reproduction. These observations have been used to suggest that improved survival may not be the selective benefit of the DR response (Adler & Bonduriansky, 2014). Instead, another hypothesis proposes that the selective benefit of the DR response is through its effect on immediate reproduction (Adler & Bonduriansky, 2014), termed the nutrient recycling hypothesis (NRH) (Regan et al., 2020). This hypothesis is based on the general finding that DR results in the inhibition of nutrient sensing pathways, e.g. TOR and IIS pathways (Adler & Bonduriansky, 2014). Inhibition of these pathways disinhibits (upregulates) nutrient recycling mechanisms such as apoptosis (James et al., 1998) and autophagy (Hansen et al., 2008; Kenyon, 2010; Fontana et al., 2010, both reviewed in Longo & Fontana, 2010). The NRH suggests that apoptosis and autophagy allow the organism to use stored nutrients from cells whilst limiting the number of cells (Adler & Bonduriansky, 2014). The individual can use available resources more efficiently, with a possible lower resource requirement for reproduction (Adler & Bonduriansky, 2014).

The NRH posits that lifespan extension is an artefact of laboratory conditions. Upregulation of apoptosis and autophagy may promote survival and limit rates of aging due to protecting against common laboratory causes of death, such as cancer or other old age pathologies (Adler & Bonduriansky, 2014; Longo & Fontana, 2010; Salomon & Rob Jackson, 2008; Spindler, 2005; Zhang & Herman, 2002). However, the limit on cell numbers and cellular growth rate may also limit the ability of individuals under DR to respond to additional stresses (Adler & Bonduriansky, 2014), with the prediction that DR would not extend lifespan in the wild (Adler & Bonduriansky, 2014). Thus, in contrast to the RRH, there is a clear prediction from the NRH that the addition of stressors, particularly injury and infection, should result in the removal or even reversal of the lifespan benefit of DR (Adler & Bonduriansky, 2014).

The effect of DR has been subject to relatively few studies in the context of injury and infection stress. In terms of injury stress, decreased calorie intake slows down wound repair in both rodents and reptiles (French et al., 2007; Hunt et al., 2012; Reed et al., 1996; Reiser et al., 1995). However, studies manipulating both overall calories and macronutrient content suggest that the main driver of the DR response, particularly in insects, is macronutrient ratio, with low protein and high carbohydrate diets leading to longer lifespans (e.g. Le Couteur et al., 2016; Kwang Pum Lee et al., 2008; Nakagawa et al., 2012; Simpson & Raubenheimer, 2009). In terms of infection stress, evidence for effects of protein to carbohydrate (P:C) ratios on proxies of survival after infection are mixed. In infected caterpillars, higher protein increases performance, measured as the product of weight gain and survival to pupation (K P Lee et al., 2006; Povey et al., 2009, 2014), and lengthens the time to death for caterpillars dying post-infection prior to pupation (Cotter et al., 2019; Wilson et al., 2020). In adult *Drosophila melanogaster,* higher protein increased survival 24 hours post-infection with bacterial infection (Kutzer et al., 2018) and higher protein as extra yeast on top of food increased number of days alive post-infection with a fungal pathogen (Le Rohellec & Le Bourg, 2009). In contrast, higher protein decreased survival measured up to 160 hours post-infection (J.-E. Lee et al., 2017), 16 days post-infection in *D. melanogaster* (Ponton et al., 2020), and decreased survival 9 days post-infection in *Bactrocera tryoni* (Dinh et al., 2019). However, to date none of these experiments have directly measured the key trait of lifetime survival. Additionally, studies often only use a small number of diets (Dinh et al., 2019; Kutzer et al., 2018; Le Rohellec & Le Bourg, 2009; J.-E. Lee et al., 2017; Ponton et al., 2020), or manipulate both P:C and calories at the same time (Kutzer et al., 2018; Le Rohellec & Le Bourg, 2009; J.-E. Lee et al., 2017), making it hard to disentangle which aspect of the diet is affecting survival with injury or infection. Furthermore, no experiments have directly compared the effect of multiple diets on lifetime survival and reproduction in control, injured and infected individuals and thus tested the alternative predictions of the current evolutionary explanations of the DR response.

Here we address this gap in our knowledge by testing the contrasting predictions of the current evolutionary explanations of the DR response by including additional stressors of injury and infection to dietary restricted *D. melanogaster.* We achieved DR by altering the P:C ratio of food (e.g. Jensen et al., 2015; Kwang Pum Lee et al., 2008) and thus throughout use the term protein restriction, although we acknowledge here this also means the associated increase in carbohydrate. We measured lifespan, reproduction, and indicators of aging, specifically the maintenance of gut integrity and climbing ability. These measures of aging are often used to track treatment specific declines in function (e.g. Grotewiel et al., 2005; Martins et al., 2018) and allows us to measure whether ageing is delayed with DR under all stress treatments. We predict that if the RRH explains DR responses, all treatments would see the usual pattern of DR, where decreasing protein increases survival up to a point and then survival declines again due to malnutrition (see review Mair & Dillin, 2008). Regardless of the stress treatment, reproduction would increase with increasing protein and ageing would be delayed with lower protein. If the NRH explains DR responses, we would expect to see that with injury and infection, the lifespan increase expected under DR would disappear and injured and infected flies would not have the usual hump shape response of lifespan to decreasing protein in the diet. In addition, infected or injured individuals would not show delayed ageing with DR. Overall, only the control group with no stress treatment would show the usual DR responses.

## METHODS

### FLY STOCKS AND MAINTENANCE CONDITIONS

We used an outbred population of *Drosophila melanogaster,* created by crossing 113 Drosophila Genetic Resource Panel (DGRP) (Mackay et al., 2012) lines in 100 pairwise crosses (consisting of two age-matched virgin females and two age-matched males from different DGRP lines; see supplementary methods) in vials containing modified Lewis food (Lewis, 1960, see Table S1, 14% protein diet). The first generation of the outcross, referred to as the Ashworth outcrossed DGRP population, was made by placing all offspring from these initial pairwise crosses in a population cage and allowing them to interbreed and lay eggs. These were collected and deposited into bottles containing Lewis food, following the method of Clancy and Kennington (2001) for maintaining *Drosophila* populations at constant densities. To generate the next generation, each month the emerged adult flies from these bottles were pooled into a population cage to lay eggs following the same method of Clancy and Kennington (2001) (more information in supplementary methods). In this way, the outcrossed population was housed in plastic bottles and outbred for 19 non-overlapping generations of complete outcrossing in 12 h light:dark cycles, at 25 °C (±1 °C) and constant humidity. Many of the original DGRP lines carry the bacterial endosymbiont Wolbachia (Mackay et al., 2012). The DGRP panel maintained the lab was cleared of Wolbachia over seven years prior to the creation of the outcrossed population.

From the 20^th^ overlapping generation of this outcrossed population, 4 μl of egg solution were placed into 20 plastic vials with modified Lewis food. After one generation, the adults were split into 50 vials, and to 60 vials from the second generation onwards. To create each generation, adults were transferred to new vials and allowed to lay eggs for two days before removal. Flies used for the experiment were offspring of the fifth generation from this protocol. The DGRP outcrossed population tested negative for common *Drosophila* laboratory viruses using primers described in Webster et al. (2015) with RT-PCR (M. A. Wallace, data not shown, March-April 2017).

### EXPERIMENTAL METHODS

Adults of the fifth generation were density controlled (10 females/vial) to minimise subsequent variation in larval densities across vials, which can affect adult life-history traits (Graves & Mueller, 1993). Mated females were allowed to lay eggs for two days and then were removed. Vials were checked daily for adult eclosion. Flies were then maintained in vials for five days after adult eclosion began to allow mating to occur after which mated female flies from over 30 of these vials were transferred into the experiment following handling under CO_2_ anaesthetisation. At this point, individual flies were singly housed on one of the ten diet treatments for the first experimental day (see below). On experimental day 2, flies from each diet treatment were assigned to one of three stress treatments: control, injury or infection (see below). There were 20 replicate flies per diet and stress treatment combination (20 individuals x 3 treatments x 10 diets = 600 flies in total).

#### DIET TREATMENTS

For the adult lifespan of each fly, flies were maintained on one of ten diets varying in protein to carbohydrate (P:C) ratio. These diets were made by altering the mass of yeast or sugar added to the modified Lewis food recipe (Lewis, 1960, Table S1). These diets were a span of P:C values (from 1:26 to 2.5:1 P:C), where protein restriction has previously been shown to extend lifespan (Kwang Pum Lee, 2015).

#### STRESS TREATMENTS

On experimental day 2, flies were exposed to one of three stress treatments: control, injury or infection. The control treatment involved handling flies under CO_2_ anaesthetisation and then transferring these to a new vial containing the relevant diet. The injury treatment involved the same protocol, however an enamelled pin was dipped in sterile LB broth and used to pierce the pleural suture under the left wing. For the infection treatment, the pin was dipped in a *Pseudomonas entomophila* bacterial broth from an overnight culture in LB at 30 °C and used to pierce the pleural suture under the left wing (following Dieppois et al., 2015; Troha & Buchon, 2019, see also Chakrabarti et al., 2012; Vodovar et al., 2005 for more information on the pathogen). To avoid lethal or negligible doses, an OD of 0.005 of *P. entomophila* culture was used for infections, as determined in a previous pilot study (J. A. Siva-Jothy, data not shown, November, 2017).

### SURVIVAL AND FECUNDITY MEASURES

Individuals were followed for life with survival scored daily. For the first two weeks of the experiment, individuals were tipped into fresh vials daily and afterwards every second day, with eggs (hatched and unhatched) counted when tipped. Any additional eggs in the vial were counted if a fly died on a day without a scheduled egg count. Diets and stress treatments were randomised across trays and trays were moved around the incubator daily to minimise microclimate effects.

### MEASURES OF PHYSIOLOGICAL AGEING

#### GUT DETERIORATION (SMURF) ASSAY

In *D. melanogaster,* and other species (Martins et al., 2018), physiological ageing is associated with increased gut permeability, which can be assessed by feeding flies food with a blue dye and observing a change in body colour if the dye leaks from the gut (Rera et al., 2011). All diets included a blue food dye following Rera et al. (2011) at a lower concentration (Table 1), to allow individuals to be scored for the “smurf” phenotype with age (Rera et al., 2011). Flies were scored as smurfs if the full body was blue, rather than just a small amount of blue in the abdomen (Rera et al., 2011).

#### NEGATIVE GEOTAXIS (NG) ASSAY

As flies age, their escape response declines and this deterioration can be measured with a negative geotaxis (NG) assay (e.g. Arking & Wells, 1990; Gargano et al., 2005; Linderman et al., 2012). NG was measured once every two weeks from week three, with a method modified from Arking and Wells (1990, see supplementary methods). Briefly, flies were individually tipped into clean vials, knocked down to the bottom and then scored for whether they climbed to 4 cm on the vial within 60 seconds (1 for passing line, 0 for not passing the line).

### STATISTICAL METHODS

The data were analysed using R software, version 3.5.2 (R Core Team, 2014) and all graphs were drawn using ggplot2 (Wickham, 2016). Diet was analysed as a continuous covariate representing the percentage of protein in the diet (Table S1) and its quadratic effect to allow for non-linear effects, while stress treatment was analysed as a categorical fixed effect. To avoid scaling errors when fitting quadratic effects, all variables were standardised to a mean of zero with a standard deviation of one. This was done separately for each test due to different sample sizes for different traits.

#### SURVIVAL

We used the R Survminer package (Kassambara & Kosinski, 2018) to graph Kaplan-Mayer curves individually for each stress treatment with diet as a factor. Our survival data did not follow the assumptions of a Cox proportional hazards model (see supplementary methods), and therefore we used an event history model where survival was analysed as a binomial trait, with each day a fly scored as a 0 for being alive and 1 for dead, following Moatt et al. (2019). We used the R package MCMCglmm (Hadfield, 2010) to model survival as a binomial variable with a categorical model. The model contained the fixed effects of stress treatment, protein content and its squared term (to model non-linear effects) and their interaction. Censored flies were included in the analysis (27 individuals, so 4.5% of the total), scoring a 0 until the day of censoring. A random effect of individual identity was included to account for repeated measures on the same individual and a random effect of experimental day was added to account for variation in survival across days. Parameter expanded priors were placed on all random effects (*V* = 1, *nu* = 1, *alpha.mu*. = 0, *alpha*. *V* = 1000). The residual variance was fixed to 1, as it is inestimable in a binomial model. The model was run for 5,200,000 iterations, with a burnin of 1,200,000 iterations and a thinning interval of 4,000 iterations to minimise autocorrelation. Autocorrelation was checked from plots of the posterior distribution of all estimates for this and all subsequent models.

We also analysed lifespan to confirm the results of the survival analysis. Lifespan, the number of days an individual survived, was analysed using a generalised linear model with MCMCglmm. Censored flies were removed from the analysis. A Poisson family error distribution was assumed and the model was run for 65,000 iterations with a thinning interval of 50 iterations and a burnin of 15,000 iterations to minimise autocorrelation. Protein content, its squared term, stress treatments and their interactions were included as fixed effects. An inverse Gamma prior was placed on the residual variance (*V* = 1 and *nu* = 0.002).

#### REPRODUCTION

Lifetime reproduction was measured as the sum of all eggs counted per female over her life. The effect of stress treatment, protein content, its squared term and their interactions were analysed using a MCMCglmm model with a Poisson error distribution. The model was run for 130,000 iterations, with a burnin of 30,000 iterations and a thinning interval of 100 iterations to minimise autocorrelation. An inverse Gamma prior was placed on the residual variance (*V* = 1 and *nu* = 0.002). To remove the effect of lifespan on reproduction, the same model with the effect of mean centered lifespan for each fly was analysed separately, except with 650,000 iterations, a burnin of 150,000 iterations and a thinning interval of 500. As an additional analysis to remove the effect of lifespan on reproduction and to compare our data with other studies using measures of early reproduction, early egg production was analysed separately. Egg counts from experimental day 2 (day after stress treatment) to day 7 were considered, as the first day egg counts were very low and were very similar across diets (Figure S1). Only individuals which lived to day 7 were considered. A MCMCglmm model with a Poisson error distribution was run with 260,000 iterations, a burnin of 60,000 iterations and a thinning interval of 200 iterations. An inverse Gamma prior was placed on the residual variance (*V* = 1 and *nu* = 0.002). The effect of stress treatment, protein content and its squared term were included in the model.

#### REPRODUCTIVE AGEING

To investigate reproductive senescence, daily egg counts were analysed using MCMCglmm with a Poisson error distribution. When egg counts changed from daily to every second day counting, all values that correspond to eggs produced over two days were divided by two and rounded down to the nearest integer. Fixed effects included stress treatment, protein content and age (in days) and their squared terms, and all interactions. Mean centred lifespan was included as a fixed effect to control for selective disappearance (Van de Pol & Verhulst, 2006) and individual ID was included as a random effect to control for repeated measures on the same individual. Models were run for 2,600,000 iterations, with a thinning interval of 1,500 and a burnin of 600,000. A parameter expanded prior was used for the random effect of individual (*V* = 1, *nu* = 1, *alpha.mu* = 0, *alpha. V* = 1000) and an inverse Gamma prior placed on the residuals (*V* = 1 and *nu* = 0.002).

#### GUT DETERIORATION (SMURF) ASSAY

A fly was scored as a smurf if it developed a non-disappearing blue body appearance (1 for smurf, 0 for no smurf) at any point during its life. This binomial variable was analysed with a categorical model using MCMCglmm. This model included the fixed effects of stress treatment, protein content, its squared term and their interactions. Models were run for 26,000,000 iterations, with a thinning interval of 20,000 and a burnin of 6,000,000. The residuals variance was fixed to 1 as explained above.

#### NEGATIVE GEOTAXIS (NG) ASSAY

We analysed the data from the negative geotaxis experiments as a binomial variable (1 for climbing 4 cm in 60 seconds, 0 for failing to do this) using a categorical family in MCMCglmm. Stress treatment, protein content and age and their squared terms, their interactions and mean centred lifespan were included as fixed effects and individual identity as a random effect. The model was run for 3,900,000 iterations, with a thinning interval of 3,000 and a burnin of 900,000. A parameter expanded prior was used for individual identity (*V* = 1, *nu* = 1, *alpha.mu* = 0, *alpha. V* = 1000) and the residual variance was fixed to 1 as explained above.

## RESULTS

### SURVIVAL AND LIFESPAN

Analysing the survival data with an event history binomial model, protein had a significant non-linear effect on survival, with survival highest on diets containing an intermediate protein level (Figure 1 and S2; Table S2; Protein^2^ = 0.48 (95% credible interval (CI) = 0.26 to 0.71), p = <0.001). Stress treatment had a significant effect on survival, with individuals exposed to infection having a greater risk of death compared to control individuals for the duration of the experiment (Table S2; Infection = 0.66 (95% CI = 0.28 to 1.10) p = 0.002). There was no significant difference between injury and control treatments (Table S2; Injury = 0.14 (95% CI = −0.32 to 0.57), p = 0.54). There was a significant interaction between protein and stress, with survival increasing more rapidly from low to intermediate protein levels for the infected treatment than for any other treatment (Figure 1 and S2; Table S2; Infection:Protein = −0.31 (95% CI = −0.57 to −0.10), p = 0.004). Survival was still maximised at relatively similar protein levels across treatments and the improvement in survival with reduced protein from very high protein levels (i.e. the classical DR response in *D. melanogaster*) did not differ across treatments (Figure 1 and S2; Table S2; Injury:Protein^2^ = −0.16 (95% CI = −0.51 to 0.18), p = 0.36; Infection:Protein^2^ = −0.01 (95% CI = −0.33 to 0.30), p = 0.99). Analysing lifespan (in days) showed very similar patterns to the binomial survival analysis (Figure S3 and S4; Table S3). Although our survival data violated the Cox proportional hazards model assumptions (see supplementary methods), the results from a Cox proportional hazards model were similar to those from the event history and lifespan models (Figure S5; Table S4).

**Figure 1:**
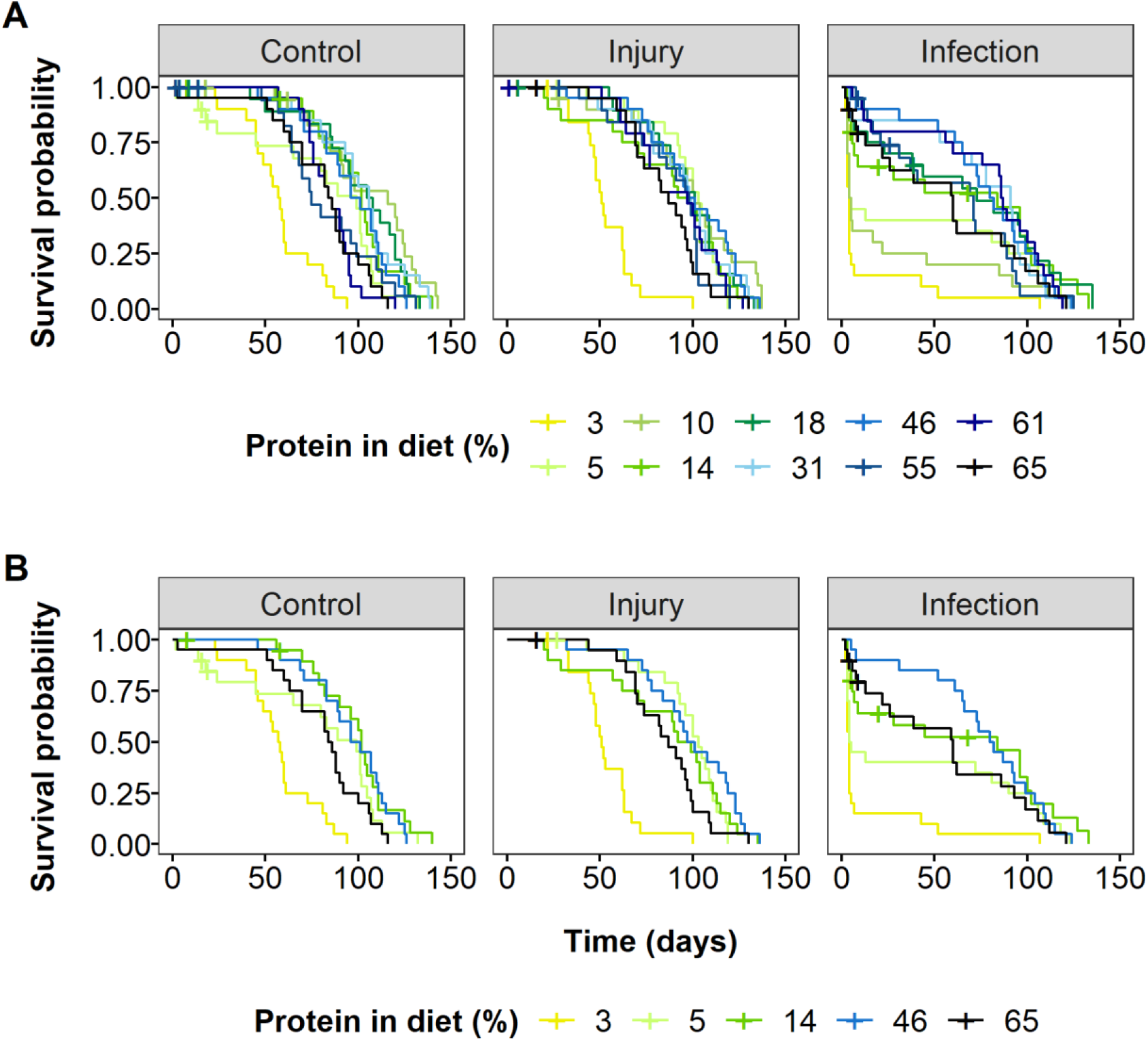
Effects of protein restriction on survival of flies infected with a bacterial pathogen (“Infection”), injured by a pinprick (“Injury”) or with no treatment (“Control”). Survival is shown as Kaplan-Meier curves for each stress treatment and protein restriction diets (A). For ease of interpretation, a subset of protein restriction diets is shown in (B) to illustrate the effects of protein restriction with low (yellow and green lines), intermediate (light blue lines) and high protein content (dark blue and black lines). Survival was maximized on intermediate protein across all stress treatments, as survival was poor on low (yellow line) and high protein diets (black line). Plus signs (+) indicate censored data points.

### REPRODUCTION

Lifetime egg production was highest at high but not the highest protein levels, with flies on low protein diets in particular producing very few eggs (Figure 2 and S6; Table S5; Protein = 1.45 (95% CI = 1.23 to 1.64), p = <0.001; Protein^2^ = −1.36 (95% CI = −−1.68 to −1.02), p = <0.001). Stress treatment had no significant effect on the lifetime number of eggs produced (Table S5; Injury = 0.19 (95% CI = −0.34 to 0.72), p = 0.49; Infection = −0.33 (95% CI = −0.90 to 0.16), p = 0.26), but there was a significant interaction between stress treatment and both protein and its squared term (Table S5; Infection:Protein = 0.47 (95% CI = 0.16 to 0.77), p = 0.01; Infection:Protein^2^ = −0.47 (95% CI = −0.93 to −0.04), p = 0.04). These interactions suggests that infected individuals had a higher linear increase in lifetime eggs with increasing protein, but this relationship was also more curved, than in either the control or injury group. Despite these significant interactions, the broad pattern of change in egg counts with changing protein level is similar across stress treatments (Figure 2).

**Figure 2:**
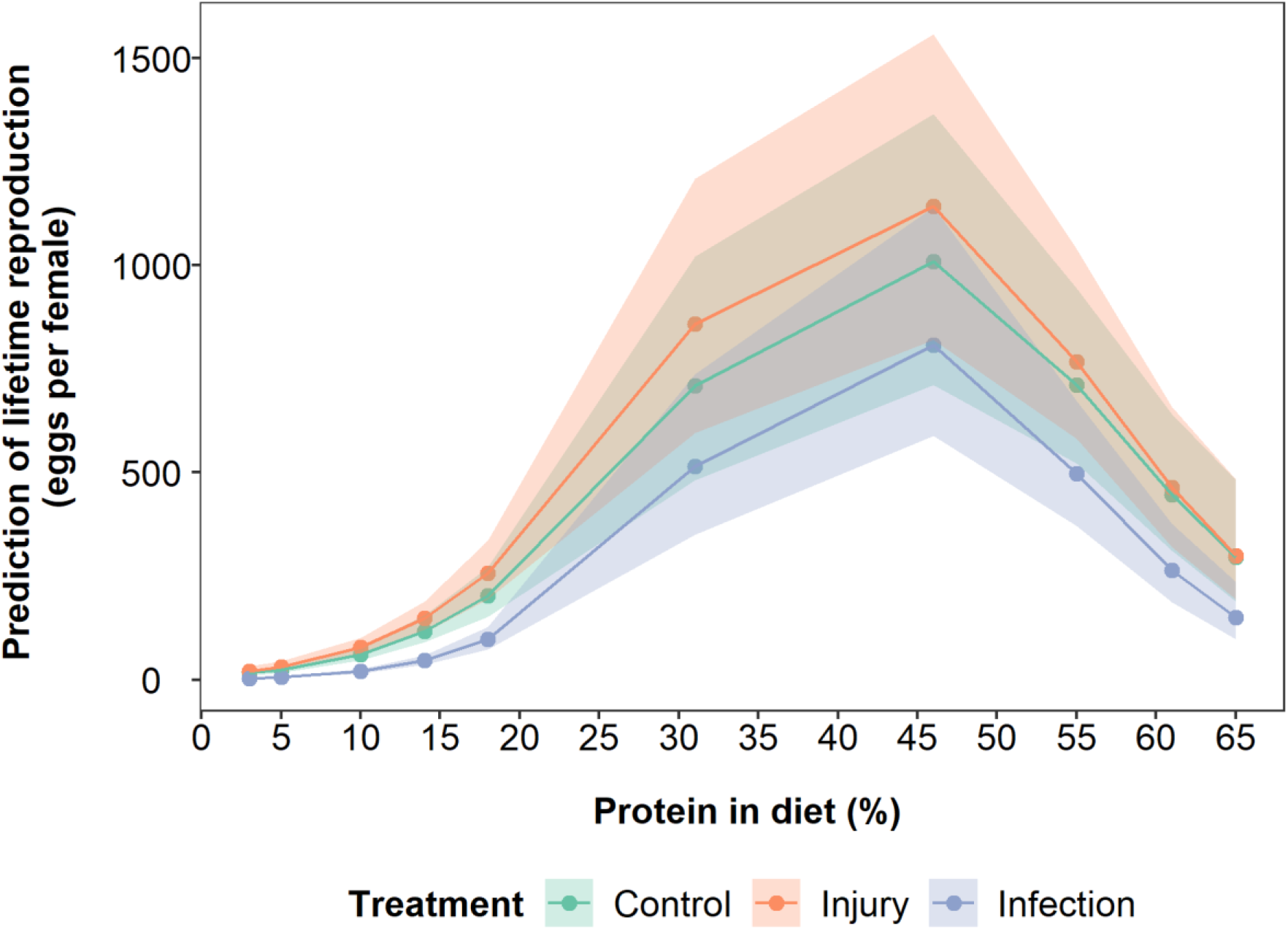
Model predictions of the effect of protein restriction on the lifetime egg production of flies infected with a bacterial pathogen (blue data points and lines), injured by a pinprick (orange data points and lines) or with no treatment (green data points and lines). Shaded areas are 95% credible intervals. Protein and protein^2^ are mean centered to standard deviation of 1.

To control for variation in lifetime egg production due to differences in lifespan, early-life egg production was also analysed. For eggs produced in the first week, excluding the first day, the patterns were similar to those of lifetime egg production (Figure S6, S7 and S8; Table S5 and S6). However, the decline in egg production at higher protein levels was reduced, such that early-life egg production plateaus after reaching a maximum at intermediate protein levels, with a slight decline at very high protein levels (Figure S8; Table S6; Protein^2^ = −0.86 (95% CI = −1.34 to −0.41), p = <0.001). There was no effect of stress treatment on early-life egg production (Figure S8; Table S6; Infection:Protein = −0.24 (95% CI = −0.74 to 0.20), p = 0.32; Infection:Protein^2^ = −0.14 (95% CI = −0.85 to 0.57), p = 0.74). Similar patterns were seen in models of lifetime egg production with mean centred lifespan included in the model (Figure S9; Table S7), suggesting that differences in lifetime reproduction between stress treatments are driven by the short lifespan of infected flies on low protein diets (Figures 2, S8 and S9). As might be expected, flies with longer lifespans had more eggs over their life than shorter-lived flies (Table S7, Lifespan = 0.93 (95% CI = 0.83 to 1.04), p = <0.001).

### AGEING

#### DAILY EGG PRODUCTION

Overall, individuals produced most eggs per day early in life, with significantly declining egg production with age (Figure 3 and S10; Table S8; Age = −0.32 (95% CI = −0.40 to −0.23), n = <0.001), but this decline was non-linear (Figure 3 and S10; Table S8; Age^2^ = −0.52 (95% CI = −0.59 to− −0.44), p = <0.001). With higher protein, individuals were able to produce significantly more eggs per day (Figure 3 and S10; Table S8; Protein = 1.31 (95% CI = 1.12 to 1.52), p = <0.001). However, at very low and high levels of protein, egg production reduced (Figure 3 and S10; Table S8; Protein^2^ = −1.5 (95% CI = −1.51 to −1.81), p = <0.001). There were numerous significant two-, and three-way interactions in the model. Overall, these interactions suggest that the curved relationship between reproduction and age is greatest for infected individuals on intermediate to high (but not the highest) protein diets (Figure 3 and S10; Table S8). Injured individuals show a similar pattern to infected individuals, but the curvature with age is generally less than for infected individuals (Figure 3 and S10; Table S8). For control individuals, again the decline in reproduction with age is steepest at higher protein levels, but not at the highest protein levels and the relationship between age and daily egg production is least curved. There was a significant effect of lifespan on daily egg production, suggesting that longer-lived individuals produced more eggs per day (Table S8; Lifespan = 0.21 (95% CI = 0.11 to 0.31), p = <0.001).

**Figure 3:**
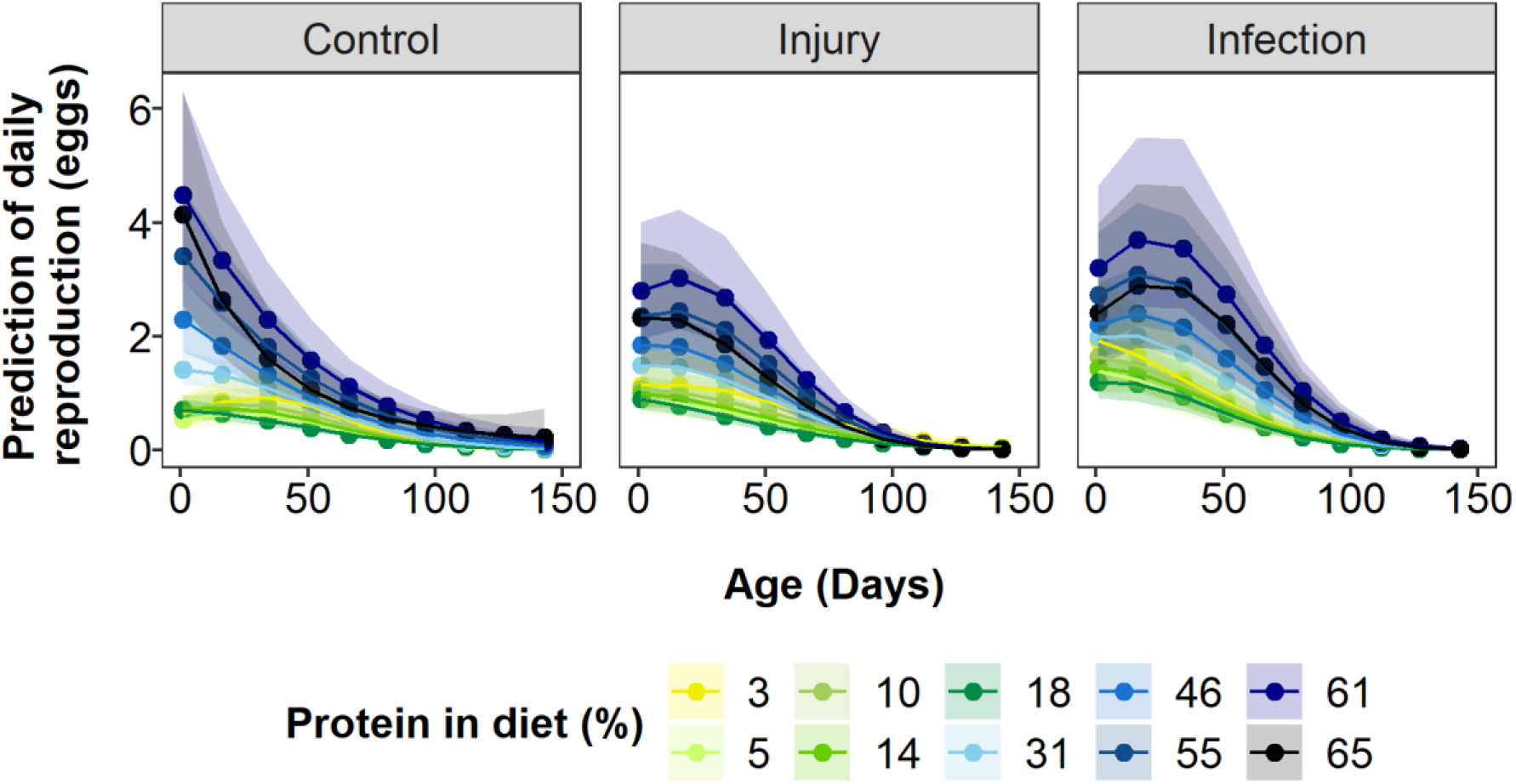
Model predictions of the effect of protein restriction and age on daily egg production of flies infected with a bacterial pathogen (“Infection”), injured by pinprick (“Injury”) or with no treatment (“Control”). Shaded areas are 95% credible intervals. Protein, protein^2^ and lifespan are mean centered to standard deviation of 1.

#### GUT DETERIORATION (SMURF) ASSAY

To assess gut integrity as a measure of ageing, flies were fed blue food and were scored as a smurf if they turned blue due to the blue food leaking from the gut. Only 11.0% of flies (63/573, excluding censored flies) became smurfs throughout the experiment, so these results should be interpreted with some caution. Diets lower in protein had more smurfs (Figure S11 and S12; Table S9; Protein = −0.75 (95% CI = −1.24 to −0.21), p = 0.004). There was a significant two-way interaction between injury treatment and protein content, where the decline in the proportion of smurfs with increasing protein content was stronger in the injury treatment than in the control treatment (Figure S12; Table S9; Injury:Protein = −1.96 (95% CI =−4.07 to −0.11), p = 0.01). There was also a significant interaction between stress treatment and the quadratic effect of protein (Table S9; Injury:Protein^2^ = −2.09 (95% CI = −4.22 to −0.47), p = 0.14; Infection:Protein^2^ = −1.73 (95% CI = −3.13 to −0.37), p = 0.01). This suggests that in infected individuals, the proportion of smurfs peaked at intermediate protein levels and then declined at both high and low protein levels. As smurfs start appearing at a later-life stage, low survival in the infected individuals on high and low protein diets may be driving this effect.

#### NEGATIVE GEOTAXIS (NG) ASSAY

By assessing escape response as a measure of ageing, protein had a significant non-linear effect on the proportion of flies passing the negative geotaxis test (Figure S13 and S14; Table S10). The likelihood of passing the test decreased with increasing protein (Table S10; Protein = −0.65 (95% CI = −1.01 to −0.32), p = <0.001), but the rate of this decline slowed at the highest protein levels (Table S10; Protein^2^ = −0.70 (95% CI = −1.21 to −0.21), p = 0.01). Overall, there were no differences between control, injured or infected flies in passing the test (Figure S13 and S14; Table S10). Older flies were less likely to pass the test (Table S10; Age = −3.57 (95% CI = −4.04 to −3.07), p = <0.001). There was an effect of selective disappearance, where longer-lived individuals passed the test at a higher rate than individuals with shorter lifespans did (Table S10, Lifespan = 0.84 (95% CI = 0.64 to 1.02), p = <0.001). Having controlled for lifespan, the proportion of flies passing the NG test declined more steeply with age on higher protein diets (Figure S14, Table S10, Protein:Age = −0.78 (95% CI = −1.06 to −0.49), p = <0.001).

## DISCUSSION

Our results provide a rare test of the predictions of two alternative evolutionary explanations for the commonly observed extension of lifespan in response to dietary restriction (DR). In particular, we tested the predictions of the nutrient recycling hypothesis (NRH) that DR will not extend lifespan with the addition of injury and infection to the usually benign laboratory environment (Adler & Bonduriansky, 2014). Alternatively, the resource reallocation hypothesis (RRH) does not make this prediction (Shanley & Kirkwood, 2000). We applied both multiple stressors and diets ranging in protein to carbohydrate (P:C) ratios to a population of outbred female *Drosophila melanogaster* to test these predictions. Our data showed that lifespan extension and delayed ageing with DR remained even with the addition of injury and infection, therefore supporting the RRH.

A small number of other studies have also considered predictions from the NRH. One tested the prediction that reproduction should decline if autophagy is inhibited under DR, but found that this was not the case in *Caenorhabditis elegans* (Travers et al., 2020). An experimental evolution study in *D. melanogaster* males hypothesised that according to the NRH, individuals under DR should be more efficient at using the available resources, and thus under long-term DR, experimental evolution lines should evolve to have higher reproductive performance and increased survival with DR (Zajitschek et al., 2016). Against their predictions for support of the NRH, there was no change in survival, although the DR selection lines did have higher reproductive performance (Zajitschek et al., 2016). One of the most direct tests of the NRH would be to investigate the effect of DR on lifespan in the wild. A recent study using wild and captive antler flies found that protein restriction lowered mortality rate even in non-laboratory conditions (Mautz et al., 2019), contradicting the suggestion of the NRH that DR would have no benefit in the wild due to higher extrinsic mortality rate and stressors (Adler & Bonduriansky, 2014). This pattern was only present in one of the two years included in the study, highlighting the need for further studies. In general it appears that the predictions of the NRH are not being met in the studies conducted to date (Adler & Bonduriansky, 2014).

Across all stress treatments, survival and lifespan were maximised at intermediate protein levels and declined at very high and low protein levels, typical of the DR response through P:C ratios (Carey et al., 2008; Kwang Pum Lee, 2015; Skorupa et al., 2008) or through other methods of DR (e.g. Bishop & Guarente, 2007; Clancy et al., 2002; K P Lee et al., 2006; Magwere et al., 2004; Pletcher et al., 2005 see also meta-analysis Nakagawa et al., 2012). In particular, survival on low protein diets was very low for infected flies, suggesting that protein is important for survival when exposed to infection. Although our use of multiple stressors and diets and monitoring lifetime survival rather than survival proxies is novel, some previous work on DR and stress treatments in insects is relevant to our results. In a study using a range of temperatures and P:C diets in *D. melanogaster,* increasing temperature both reduced lifespan and reduced the decrease in lifespan with higher protein, although there still appeared to be a slight decrease in lifespan on the highest protein diets (Kim et al., 2020). In terms of infection, higher protein increased larval performance (a product of survival and weight gain) of infected caterpillars prior to pupation in *Spodoptera littoralis* (K P Lee et al., 2006) and *Spodoptera exempta* (Povey et al., 2009, 2014). Also, for *Spodoptera littoralis* caterpillars which died post-infection, time to death is lengthened on higher protein diets (Cotter et al., 2019; Wilson et al., 2020). Similar protein effects have been found in *D. melanogaster* even though survival was only measured for a short time post-infection. With a fungal infection, the addition of live yeast on top of food was found to increase the number of days alive post-infection (Le Rohellec & Le Bourg, 2009). In the same host-pathogen study as used here, yeast restriction reduced *D. melanogaster* survival 24 hours post-infection with *Pseudomonas entomophila*, but there was no effect of yeast restriction on survival 24 hours post-infection with *Lactococcus lactis* (Kutzer et al., 2018). It should be noted that this study only changed the amount of yeast and therefore includes both calorie and protein restriction. Together these results suggest that, although increased protein may improve survival under exposure to a stressor, there may be an optimal level of protein above which survival is reduced.

There are also a small number of recent studies that show the opposite pattern to our results, where infected individuals on lower protein diets have improved performance (Dinh et al., 2019; J.-E. Lee et al., 2017; Ponton et al., 2020). Several methodological differences make direct comparison difficult. These studies only use two or three diets, with one using liquid diets (Dinh et al., 2019), with much lower protein to carbohydrate ratios to the ones used in our study. In the one study that did use diets with comparable protein levels to ours (one diet at 52% protein), survival on this diet for 16 days post treatments in all treatments was very low (Ponton et al., 2020). In comparison, this diet is similar to the diet with the highest survival in our study. Other than methodological differences, the particular host-parasite system used across studies may be driving these differences. In a meta-analysis of the effect of host nutrition on pathogen virulence, nutrition quality or quantity did not have a consistent effect on the host performance, with both positive or negative effects on virulence depending on the system (Pike et al., 2019). To understand what effects protein has on short-term infection outcomes, many different measures of the host or pathogen have been studied. For example, higher protein has been found to decrease (Kutzer & Armitage, 2016; Wilson et al., 2020), increase (Dinh et al., 2019; J.-E. Lee et al., 2017), or have no effect on bacterial growth (Kutzer et al., 2018) post-infection. A measure of the immune response, the production of antimicrobial peptides, has either increased (K P Lee et al., 2006; Povey et al., 2009, 2014) or decreased (Ponton et al., 2020) on higher protein post-infection. In caterpillars, higher protein has been seen to increase the functional immune response post-infection (Cotter et al., 2019) and increase the amount of protein in the haemolymph (Cotter et al., 2010; Povey et al., 2009), which might be affecting the growth of bacteria (Wilson et al., 2020). To understand why lower protein lowers survival in the infected flies in our host-pathogen system in comparison to other studies better, further work on measures of the host response and pathogen are needed.

Injured flies in our study also showed a lifespan benefit in response to modest protein restriction. Surprisingly, injury had no measurable effect on survival. In *D. melanogaster,* there are conflicting results of injury effects, potentially due to the methods of applying injury. Injury to the thorax has been seen to increase lifespan in young male flies but this effect was absent in young female or older flies (Henten et al., 2016). In contrast, injury by removing leg parts increased mortality rates in the first 28 days post-injury in male but not female flies (Sepulveda et al., 2008). Both studies housed flies in groups, limiting direct comparisons to our results. In terms of diet effects on injury, injured *D. melanogaster* have been found to have similar survival to control flies 16 days post-injury but only if they were on the lowest protein diet (4% protein) of the three tested (Ponton et al., 2020). In contrast, protein had no statistically significant effect on performance prior to pupation for injured caterpillars (Povey et al., 2009). As injury in *D. melanogaster* activates immune response pathways (e.g. Agaisse et al., 2003; Hoffmann & Reichhart, 2002), indicating it is a stressor to the flies, understanding why flies on different diets do not show lifespan differences if injured is an interesting area of future research.

Although the pattern of a tent-shaped response of survival and lifespan to increasing levels of protein restriction seen here is typical of many other studies (Carey et al., 2008; Kim et al., 2020; Kwang Pum Lee, 2015; Skorupa et al., 2008), it does contrast with recent studies suggesting lifespan is maximised on diets with very low protein:carbohydrate (e.g. mice (Solon-Biet et al., 2014), crickets (Harrison et al., 2014; Maklakov et al., 2008), *B. tryoni* (Fanson et al., 2009, 2012), and *D. melanogaster* (Jensen et al., 2015; Kwang Pum Lee et al., 2008). These studies use a nutritional geometry approach where a very large number of diets that vary in both calories and macronutrient ratio are used in order to separate the effects of these two variables. One reason our results may differ is due to difference in the delivery of the diets. Most studies using nutritional geometry in *D. melanogaster* have used liquid diets that allow fine scale measures of intake, but result in very low survival rates across all diets (Jensen et al., 2015; Kwang Pum Lee et al., 2008). One nutritional geometry study using *D. melanogaster* with solid diets found that lifespan was maximised on intermediate protein and lifespan was overall greater than in the liquid diet results (Skorupa et al., 2008). A study using eight solid diets in male *D. melanogaster* found that low P:C ratio diets maximised lifespan, but that the lowest P:C ratio decreased lifespan (Bruce et al., 2013). A study using both male and female *D. melanogaster* found that low P:C ratios with solid food increased lifespan in females, but that this response was curved and the longest lifespan was at 1:4 P:C (Kim et al., 2020). This suggests that diet delivery may have effects on survival, at least in *D. melanogaster*. In comparison, *B. tryoni* have longer lifespans on liquid diets (Fanson et al., 2009, 2012) compared to *D. melanogaster* liquid diets. Other studies in crickets (Harrison et al., 2014; Maklakov et al., 2008) or mice (Solon-Biet et al., 2014) use solid diets, so more work is needed to understand the causes of the differences between studies.

Lifetime reproduction was maximised at intermediate protein levels, although at a slightly higher protein level than lifespan, a result which has been seen in other studies (Fanson et al., 2009; Harrison et al., 2014; Jensen et al., 2015; Kwang Pum Lee et al., 2008; Maklakov et al., 2008). However, some studies have shown that lifetime egg production is maximised at the highest protein level (Carey et al., 2008; Moatt et al., 2019). Patterns in lifetime reproduction could be driven by differences in lifespan rather than reproductive rate, especially as longer-lived flies have been seen to lay more eggs (Jensen et al., 2015). By incorporating lifespan into our model of lifetime egg production and by analysing early-life egg production, we found that egg production still peaked at intermediate protein levels, but with higher protein, the decline in egg production was not as steep as with lifetime eggs. Similarly, measures of egg laying rate have been found to peak at intermediate protein diets in other studies (Fanson et al., 2009, 2012; Harrison et al., 2014; Jensen et al., 2015; Kim et al., 2020; Kwang Pum Lee et al., 2008; Maklakov et al., 2008) as with early-life reproduction (Rapkin et al., 2018). However, early-life egg production has also been seen to peak at the highest protein diet (Kwang Pum Lee, 2015; Skorupa et al., 2008). These inconsistencies may be due to the slight differences in diet composition used in these studies, for example by using chemically defined diets (Kwang Pum Lee, 2015) or by not including cornmeal in the diets (Skorupa et al., 2008). However, the broad pattern that reproduction is maximised on higher protein diets than lifespan appears consistent across studies.

In terms of stress treatment and reproduction, across control, injury and infection treatments, we saw the same patterns of highest egg counts on intermediate protein. Infected flies overall produced fewer eggs in comparison to the control or injured flies, as seen in many studies focusing on the reproduction-immunity trade off (reviewed in Schwenke et al., 2016). If lifespan was accounted for in the lifetime reproduction models, or only early-life reproduction was considered, there was no difference in reproduction between the stress treatments. This suggests that the pattern of lower lifetime reproduction in infected flies is most likely due to infected flies having shorter lifespans. Similar to our results, in *D. melanogaster* yeast restriction had a larger effect on early-life egg production than infection (Kutzer et al., 2018; Kutzer & Armitage, 2016). Contrary to our results, limited availability of food and an immune challenge with dead bacteria reduced egg production rate in early-life in crickets (Stahlschmidt et al., 2013). Injury in the form of leg removal reduced egg production in the first 10 days of *D. melanogaster* housed in groups (Sepulveda et al., 2008). Additionally, oral infection with *Pseudomonas aeruginosa* increased early-life egg production but only on higher protein diets (Hudson et al., 2019). Using a range of temperatures, egg production rate in early-life in *D. melanogaster* was reduced at high and low temperatures, however intermediate to high protein diets had the highest egg production rates at intermediate temperatures (Kim et al., 2020). Higher temperatures increased egg production rate at higher protein diets (Kim et al., 2020). Therefore, the methods of stress response or the particular host-pathogen system may have an effect on the response of host reproduction on different diets.

The patterns of reproductive ageing involved complex interactions between diet and stress treatment. Broadly, there were similar patterns of ageing across treatments and diets, with an increase in egg production early in the experiment, followed by a peak and then diminishing egg numbers, as seen in other experiments (Carey et al., 2008; Le Rohellec & Le Bourg, 2009). These peaks were higher for the high protein diets (but not necessarily the highest), most likely due to the requirement of protein for egg production (Mirth et al., 2019; Wheeler, 1996). Diets with low protein levels (e.g. 3 to 18% protein) had the slowest rate of decline in egg production with age. This could simply be a result of high protein diets having much higher egg production earlier in life and thus a greater potential decline than low protein diets. It is notable that high protein diets decline rapidly in egg production early in life before the rate of decline reduces to that of lower protein diets later in life, suggesting there is an initially higher rate of ageing on higher protein diets. Additionally, the control flies have a more linear decline in egg laying, suggesting that injury and infection might slightly delay egg production. Previous studies have also found ageing in female reproduction was quicker on higher protein diets (Jensen et al., 2015; Moatt et al., 2019). Without directly testing for ageing in egg production, similar patterns of quicker declines in egg production on higher protein and calorie diets have been seen in the tephritid fruit fly (Carey et al., 2008) and in *D. melanogaster* supplemented with live yeast on top of food (Le Rohellec & Le Bourg, 2009). One study in crickets found no significant relationship between ageing in egg laying and protein in diet in females (Maklakov et al., 2009). Overall, these similarities across studies suggest diet interacts with reproductive ageing in a broadly similar way across species.

Other than ageing in reproduction, we also investigated ageing in traits that are not implicated in the survival-reproduction trade-off, as delayed ageing is a known DR response (e.g. Ingram et al., 1987; Le Rohellec & Le Bourg, 2009; Mattson et al., 2001; Regan et al., 2016; Rera et al., 2012). Measuring escape response across lifespan, we used the negative geotaxis (NG) assay, and found that ageing was delayed on lower protein diets, as has been found in another study limiting the addition of live yeast on food (Le Rohellec & Le Bourg, 2009). This is also consistent with the pattern that DR animals have an increased activity pattern (Duffy et al., 1997). A study using two genetic backgrounds of *D. melanogaster* found the effect of DR on NG was inconsistent across genotype (Bhandari et al., 2007), however they simultaneously manipulated calories and nutrient composition making direct comparison to our results difficult. We did not see effects of stress treatment on NG, in contrast to a study where infection reduced the NG response in one of two tested *D. melanogaster* genetic backgrounds (Linderman et al., 2012). These differences suggest there may be dissimilarities in the response to DR depending on the genetic background. Given the flies used in our study are genetically heterogeneous, the patterns we observe should be representative of the average genotype in this population.

We also measured the loss of gut integrity of flies with age using a smurf assay, which has been found to be more common in flies on unrestricted diets (Regan et al., 2016; Rera et al., 2012). Unexpectedly, we saw higher numbers of smurfs appearing at lower protein diets in the control and injury treatments, whilst in the infected treatment the number of smurfs was highest at intermediate protein levels. This is surprising in the infected flies as *Pseudomonas entomophila* infection is known to disrupt the gut (Chakrabarti et al., 2012; Dieppois et al., 2015). One explanation for the patterns we observe in the control and injury treatment is that the lowest protein diets used in this study are particularly low and therefore may represent malnourished conditions, leading to an increase in the number of smurfs. Nonetheless, we would still expect to see a reduction in the number of smurfs at intermediate protein levels. In addition, for infected flies, the high mortality at high and low protein levels may result in flies dying before reaching the age where smurfs start appearing. The major problem with interpreting these patterns in our experiment is the very low number of smurfs that appeared, meaning these patterns may not be robust. Additionally, we analysed the smurf trait as a binary variable, however it has been found to be a continuous trait, where at some point all individuals develop the trait (Martins et al., 2018). Therefore, by measuring the phenotype as a binary trait with only clear smurfs counted, we may have been missed some more subtle patterns. More work is required to understand how the relationship between protein restriction and the appearance of smurfs varies with exposure to injury and infection.

Overall, the addition of injury and infection did not remove the lifespan benefit of protein restriction or the delay in reproductive ageing. Our study therefore provides no evidence to support the nutrient recycling hypothesis of the lifespan response to dietary restriction. Even though there were minor differences between the stress treatments in the relationship between protein content of the diet and survival, the major pattern of survival being maximised at intermediate protein levels was maintained across stress treatments. With infection, survival was particularly poor on the lowest protein diets, whilst in the other treatment groups this difference was not as dramatic. The explanation for this pattern requires further investigation. Our results and those of other studies suggest that the resource reallocation hypothesis remains the best-supported evolutionary explanation for the lifespan benefit of dietary restriction.

## ACKNOWLEDGEMENTS

This work was supported by the Biotechnology and Biological Sciences Research Council (BBSRC) grant number BB/M010996/1. We thank Joshua Moatt for helpful advice and comments on the data analysis and write-up, Jonathon Siva-Jothy for help with the bacterial work, Katy Monteith for laboratory training and practical advice, and Megan Wallace for testing the DGRP outcross for the presence of common fly viruses.

## CONFLICTS OF INTEREST

The authors declare no conflicts of interest.

## AUTHOR CONTRIBUTIONS

ES, PV and CW designed the experiment. ES and CM did the study, with ES completing data 617 collection for the full experiment. FMW and KM created and maintained the fly population 618 used in the study and wrote the supplementary methods part for the creation of the fly 619 population. ES analysed the data with help from CW. ES wrote the paper with help from CW, 620 PV, FMW and KM.

## Notes

### Competing Interest Statement

The authors have declared no competing interest.

