## Supplement for "Testing evolutionary explanations for the lifespan benefit of dietary restriction in *Drosophila melanogaster*"

*SUPPLEMENTARY METHODS:*

**ASHWORTH OUTCROSSED DGRP POPULATION*:* A genetically diverse laboratory population resource for Drosophila melanogaster studies**

The Ashworth outcrossed DGRP population was founded by Fergal M. Waldron on 15/10/14, and is derived from 113 inbred DGRP lines sampled from a wild population in Raleigh, NC, USA (Mackay et al., 2012). The population was maintained and brought through subsequent generations of outcrossing by Fergal M. Waldron until 1/10/2015, and from 29/10/2015 by Katy Monteith.

The Ashworth outcrossed DGRP population was founded with genetic contributions from the following 113 DGRP lines; RAL-28, RAL-31, RAL-48, RAL-49, RAL-57, RAL-59, RAL-69, RAL-75, RAL-83, RAL-91, RAL-93, RAL-101, RAL-129, RAL-138, RAL-149, RAL-153, RAL-158, RAL-189, RAL-195, RAL-208, RAL-217, RAL-228, RAL-237, RAL-239, RAL-280, RAL-287, RAL-288, RAL-301, RAL-303, RAL-304, RAL-306, RAL-309, RAL-310, RAL-317, RAL-321, RAL-324, RAL-348, RAL-350, RAL-352, RAL-354, RAL-358, RAL-360, RAL-361, RAL-365, RAL-366, RAL-373, RAL-375, RAL-377, RAL-379, RAL-380, RAL-381, RAL-382, RAL-386, RAL-390, RAL-392, RAL-395, RAL-397, RAL-399, RAL-405, RAL-406, RAL-409, RAL-426, RAL-427, RAL-437, RAL-439, RAL-443, RAL-486, RAL-491, RAL-492, RAL-502, RAL-508, RAL-509, RAL-517, RAL-528, RAL-530, RAL-535, RAL-555, RAL-559, RAL-563, RAL-566, RAL-575, RAL-584, RAL-589, RAL-627, RAL-630, RAL-634, RAL-703, RAL-712, RAL-716, RAL-732, RAL-765, RAL-774, RAL-776, RAL-786, RAL-796, RAL-805, RAL-808, RAL-818, RAL-820, RAL-821, RAL-822, RAL-832, RAL-852, RAL-853, RAL-855, RAL-859, RAL-879, RAL-882, RAL-884, RAL-897, RAL-907, RAL-908, RAL-913.

To maximise the genetic contribution of each of the founder lines to the final outcrossed population, initial pairwise crosses between randomly selected population founder lines were carried out. The offspring from these pairwise crosses were then pooled into a population cage for the 1$^{st}$ generation of outcrossing. Whilst a minimum of 57 pairwise crosses would encompass inclusion of all 113 founder lines, 100 pairwise crosses were carried out as a precautionary measure against a number of crosses failing to produce offspring (an upper limit of 100 was dictated by feasibility). Pairwise crosses were set up using two virgin females crossed to two males. All virgin females and males were age-matched controlled (1-6 days old when crosses were set up). Pairwise crosses were set up in standard Lewis medium containing vials and placed at 25°C for 5 days after which adults were removed.

For pairwise crosses, DGRP outcrossed population founder lines were randomly selected to contribute females or males for the following 100 crosses (scheme is "2 virgin females from line" x "2 males from line"): RAL-390 x RAL-381, RAL-280 x RAL-301, RAL-913 x RAL-365, RAL-796 x RAL-712, RAL-589 x RAL-49, RAL-350 x RAL-382, RAL-853 x RAL-158, RAL-288 x RAL-855, RAL-49 x RAL-366, RAL-303 x RAL-908, RAL-101 x RAL-303, RAL-712 x RAL-426, RAL-321 x RAL-732, RAL-377 x RAL-101, RAL-380 x RAL-879, RAL-820 x RAL-324, RAL-882 x RAL-535, RAL-439 x RAL-634, RAL-83 x RAL-409, RAL-28 x RAL-75, RAL-409 x RAL-832, RAL-879 x RAL-237, RAL-237 x RAL-239, RAL-443 x RAL-776, RAL-908 x RAL-627, RAL-59 x RAL-584, RAL-365 x RAL-796, RAL-634 x RAL-405, RAL-392 x RAL-852, RAL-129 x RAL-350, RAL-317 x RAL-306, RAL-427 x RAL-528, RAL-373 x RAL-502, RAL-386 x RAL-28, RAL-304 x RAL-392, RAL-774 x RAL-555, RAL-306 x RAL-386, RAL-310 x RAL-309, RAL-832 x RAL-287, RAL-405 x RAL-280, RAL-57 x RAL-774, RAL-627 x RAL-228, RAL-397 x RAL-821, RAL-348 x RAL-492, RAL-437 x RAL-443, RAL-91 x RAL-31, RAL-352 x RAL-575, RAL-301 x RAL-390, RAL-48 x RAL-897, RAL-575 x RAL-808, RAL-426 x RAL-373, RAL-375 x RAL-195, RAL-31 x RAL-59, RAL-897 x RAL-310, RAL-239 x RAL-486, RAL-287 x RAL-805, RAL-584 x RAL-765, RAL-381 x RAL-149, RAL-93 x RAL-703, RAL-379 x RAL-517, RAL-821 x RAL-630, RAL-189 x RAL-853, RAL-399 x RAL-360, RAL-907 x RAL-217, RAL-535 x RAL-786, RAL-195 x RAL-395, RAL-852 x RAL-913, RAL-502 x RAL-818, RAL-361 x RAL-375, RAL-138 x RAL-491, RAL-808 x RAL-93, RAL-517 x RAL-208, RAL-153 x RAL-189, RAL-149 x RAL-352, RAL-732 x RAL-509, RAL-818 x RAL-563, RAL-630 x RAL-57, RAL-395 x RAL-380, RAL-358 x RAL-822, RAL-765 x RAL-406, RAL-703 x RAL-859, RAL-406 x RAL-153, RAL-508 x RAL-379, RAL-716 x RAL-427, RAL-509 x RAL-358, RAL-555 x RAL-48, RAL-360 x RAL-321, RAL-786 x RAL-69, RAL-855 x RAL-354, RAL-559 x RAL-437, RAL-563 x RAL-361, RAL-158 x RAL-559, RAL-805 x RAL-884, RAL-208 x RAL-566, RAL-492 x RAL-397, RAL-382 x RAL-399, RAL-75 x RAL-508, RAL-884 x RAL-138, RAL-530 x RAL-83, RAL-69 x RAL-348. Offspring from pairwise crosses were collected 28 days after parents were removed and pooled into a large *Drosophila* population cage, for the 1$^{st}$ generation of outcrossing and subsequent embryo collection.

For this, and each subsequent generation of outcrossing, the outcrossed DGRP population is maintained employing a method used to maintain constant larval densities (223 ± 14.3 (95% CI)) in stock bottles (Clancy & Kennington, 2001). Briefly, this method involves populating a large *Drosophila* cage with thousands of flies on the day 1, providing these with fruit juice (grape/apple) agar plates for embryo laying. After a 24 hr habituation period, agar plates are replaced (day 2). On the day 3, agar plates are recovered and embryos are collected from the surface. Using PBS and a brush, concentrated egg/PBS solutions are prepared, and these are squirted on the surface of Lewis media in bottles. This process is typically carried out every 20-25 days. The outcrossed DGRP populations is maintained at a density of 20-25 bottles (20 bottles maintains the population at >4000 individuals).

Below is a table outlining details/timescale of the outcrossed DGRP population foundation and maintenance (Apr ’14 – Nov ’16).

| Date | Action | Temperature (°C) | Number of bottles | Worker |
|---|---|---|---|---|
| 18/10/2014 | Tipped DGRP onto new food, and placed at 18C for virgin collection in mid-Oct | 18 | NA | FMW/WHP |
| 07/10/2014 | Tipped out adult DGRPs from vials for subsequent virgin collection | 25 | NA | FMW |
| 8/102014 - 14/10/2014 | Virgins and males collected | 25 | NA | FMW |
| 15/10/2014 | Pairwise crosses (n=100) set up (virgin male and female flies from 113 lines) | 25 | NA | FMW |
| 20/10/2014 | Pairwise crosses parents removed from vials | 25 | NA | FMW |
| 19/11/2014 | Egg squirts from offspring of pairwise crosses: 1st | 25 | 30 | FMW |

| | | | | |
|---|---|---|---|---|
| | generation of complete outcrossing | | | |
| 18/12/2014 | Tipped flies from old bottles into fresh bottles | 25 | 30 | FMW |
| 30/01/2015 | Egg squirts: 2nd generation of complete outcrossing | 25 | 30 | FMW |
| 06/03/2015 | Tipped flies from old bottles into fresh bottles | 25 | 30 | FMW |
| 09/04/2015 | Egg squirts: 3rd generation of complete outcrossing | 25 | 25 | FMW |
| 19/05/2015 | Egg squirts: 4th generation of complete outcrossing | 25 | 21 | FMW |
| 19/06/2015 | Egg squirts: 5th generation of complete outcrossing | 25 | 25 | FMW |
| 30/07/2015 | Egg squirts: 6th generation of complete outcrossing | 25 | 25 | FMW |
| 01/10/2015 | Egg squirts: 7th generation of complete outcrossing | 25 | 25 | FMW |
| 29/10/2015 | Egg squirts: 8th generation of complete outcrossing | 25 | 20 | KMM |
| 02/12/2015 | Tipped flies from old bottles into fresh bottles | 25 | 20 | KMM |
| 06/01/2016 | Egg squirts: 9th generation of complete outcrossing | 25 | 20 | KMM |
| 10/02/2016 | Egg squirts: 10th generation of complete outcrossing | 25 | 20 | KMM |
| 22/02/2016 | Tipped flies from old bottles into fresh bottles | 25 | 20 | KMM |

| | | | | |
|---|---|---|---|---|
| 25/03/2016 | Egg squirts: 11th generation of complete outcrossing | 25 | 24 | KMM |
| 26/04/2016 | Egg squirts: 12th generation of complete outcrossing | 25 | 22 | KMM |
| 18/05/2016 | Egg squirts: 13th generation of complete outcrossing | 25 | 20 | KMM |
| 14/06/2016 | Egg squirts: 14th generation of complete outcrossing | 25 | 22 | KMM |
| 14/07/2016 | Egg squirts: 15th generation of complete outcrossing | 25 | 22 | KMM |
| 10/08/2016 | Egg squirts: 16th generation of complete outcrossing | 25 | 25 | KMM |
| 07/09/2016 | Egg squirts: 17th generation of complete outcrossing | 25 | 25 | KMM |
| 04/10/2016 | Egg squirts: 18th generation of complete outcrossing | 25 | 25 | KMM |
| 02/11/2016 | Egg squirts: 19th generation of complete outcrossing | 25 | 25 | KMM |
| 30/11/2016 | Egg squirts: 20th generation of complete outcrossing | 25 | 25 | KMM |

### DIETS:

**Table S1**: Ten diets and their corresponding P:C ratios with additional information of each added ingredient. The standard modified Lewis food (Lewis, 1960) and associated P:C ratio is in bold. One of the main differences to the original Lewis food recipe is the replacement of dextrose and sucrose with brown sugar in our diets (Lewis, 1960). The P:C ratios (rounded to the nearest whole number) incorporate the protein and carbohydrate contributed by maize. Yeast and sugar are roughly isocaloric, so P:C ratios can be altered without altering the energy content of the diet by replacing yeast with sugar (Mair et al., 2005). Two baseline ratios were made with no addition of yeast (2.5:1) or sugar (1:26). All the diets were dyed using a food dye (brilliant blue FCF E133).

| P:C ratio | % Protein | Yeast (g) | Sugar (g) | Maize (g) | | | Agar (g) | Nipagin (ml) | Food dye (g) | $dH_2O$ (l) |
|---|---|---|---|---|---|---|---|---|---|---|
| | | | | Total | Of which carbohydrate | Of which protein | | | | |
| 1:26 | 3 | 0.0 | 675.0 | 415 | 290.5 | 37.8 | 41.2 | 90 | 3 | 6 |
| 1:16 | 5 | 21.3 | 653.7 | 415 | 290.5 | 37.8 | 41.2 | 90 | 3 | 6 |
| 1:8 | 10 | 73.7 | 601.3 | 415 | 290.5 | 37.8 | 41.2 | 90 | 3 | 6 |
| **1:6** | **14** | **112.5** | **562.5** | **415** | **290.5** | **37.8** | **41.2** | **90** | **3** | **6** |
| 1:4 | 18 | 162.9 | 512.1 | 415 | 290.5 | 37.8 | 41.2 | 90 | 3 | 6 |
| 1:2 | 31 | 296.7 | 378.3 | 415 | 290.5 | 37.8 | 41.2 | 90 | 3 | 6 |
| 1:1 | 46 | 463.9 | 211.1 | 415 | 290.5 | 37.8 | 41.2 | 90 | 3 | 6 |
| 1.5:1 | 55 | 564.2 | 110.8 | 415 | 290.5 | 37.8 | 41.2 | 90 | 3 | 6 |
| 2:1 | 61 | 631.1 | 43.9 | 415 | 290.5 | 37.8 | 41.2 | 90 | 3 | 6 |
| 2.5:1 | 65 | 675.0 | 0.0 | 415 | 290.5 | 37.8 | 41.2 | 90 | 3 | 6 |

### NEGATIVE GEOTAXIS (NG) ASSAY:

This assay quantifies the climbing response of flies in terms of distance or speed, following Arking and Wells (1990). A rubber band was tied 4 cm from the bottom around an empty vial. After the fly was tipped into this vial and blocked with a cotton bud, the vial was tapped down three times on a corkboard. The timer was started on the last tap and stopped once the fly fully crossed the line. After the test, the fly was transferred to a new food vial. An upper limit of 60 seconds was set as some flies did not climb or cross the line. One vial was used per fly to avoid confounding effects of reusing vials (Nichols et al., 2012) or possible spread of infection. Due to time of day effects (Gargano et al., 2005), testing order was reversed each week. If the fly did not touch the bottom of the vial, or if the timer was stopped incorrectly, a second trial was completed. Due to the number of failed tests where the fly did not cross the line (43% of 5,117 tests), negative geotaxis scores were analysed as a binomial variable for passing (1) or failing (0) the test in 60 seconds.

### COX PROPORTIONAL MODEL:

We analysed the survival data with a Cox proportional hazards model using the R Survival package (Therneau, 2015). The model included protein content, its squared term, stress treatments and their interactions as fixed effects. The assumptions of a Cox proportional hazards model were violated (Therneau, 2015, cox.zph function global term Chi squared = 95.26, p = <0.001). Predicted risk ratios for each diet and stress treatment were calculated using the predict function for the Cox proportional hazards model.

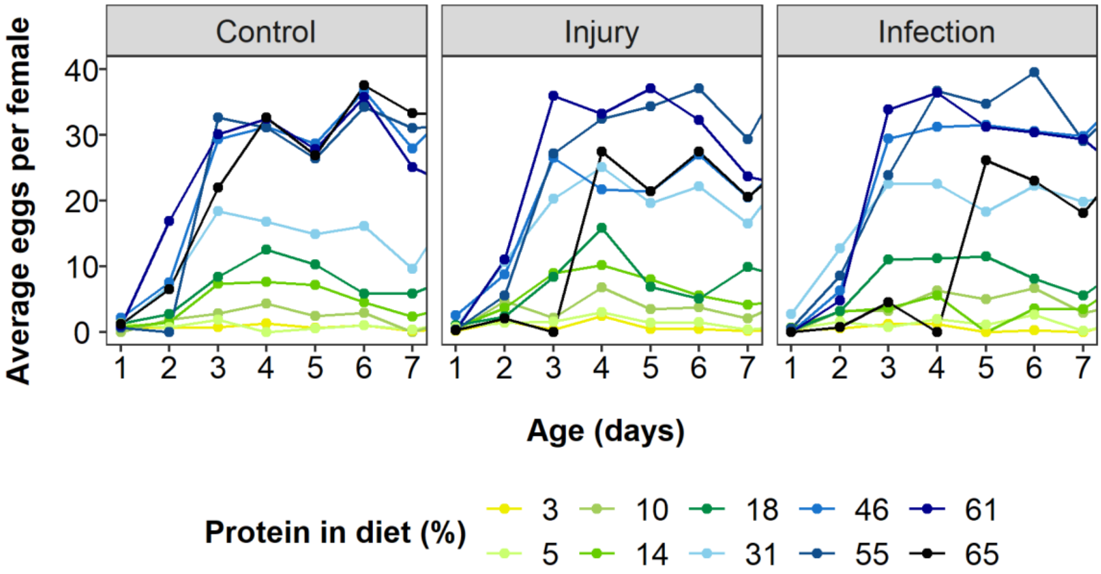


**Figure S1:** Average eggs per day produced in the first week for each protein restriction diet of flies infected with a bacterial pathogen ("Infection"), injured by a pinprick ("Injury") or with no treatment ("Control").

*SUPPLEMENTARY GRAPHS AND TABLES:*

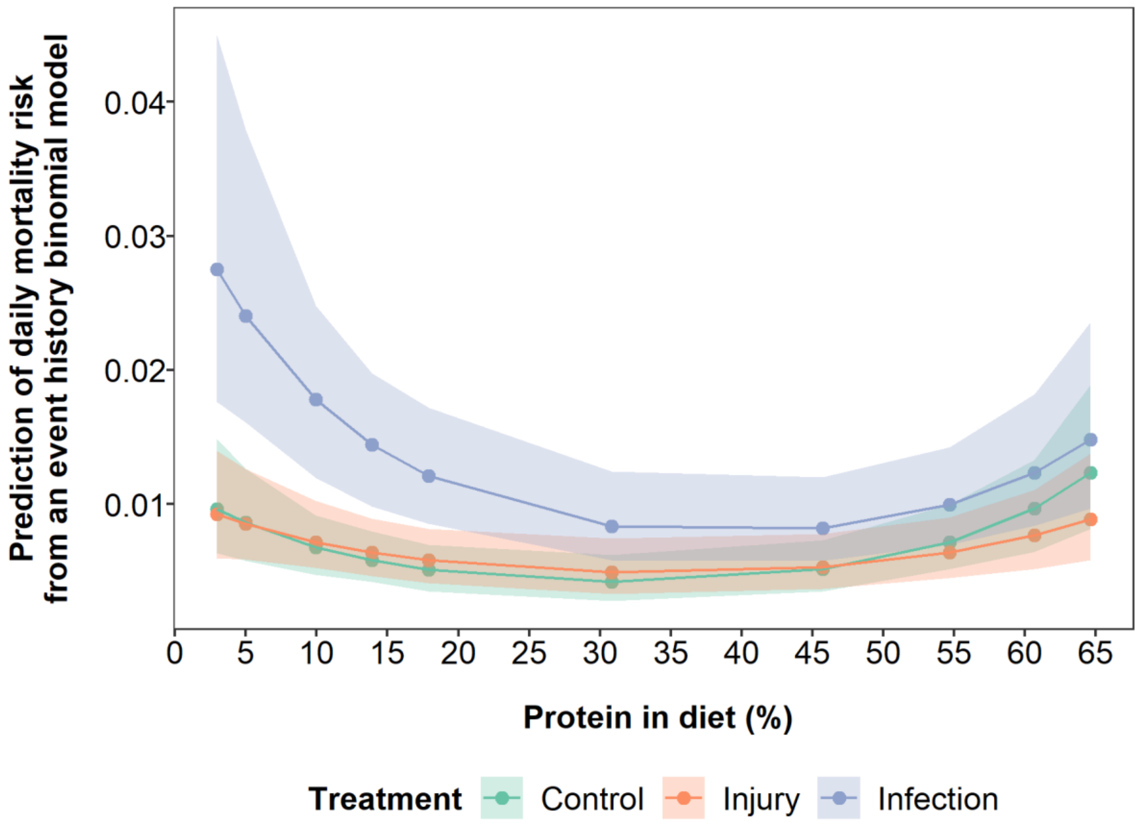


**Figure S2:** Model predictions from an event history binomial model for the effect of protein restriction on mortality risk per day of flies infected with a bacterial pathogen (blue data points and lines), injured by a pinprick (orange data points and lines) or with no treatment (green data points and lines). In the binomial model, for each day each fly was coded as 0 for alive and 1 for dead. Protein and protein$^2$ are mean centered to standard deviation of 1. Shaded areas are 95% credible intervals.

**Table S2:** Model summary of effects of protein restriction and stress treatments on mortality risk per day from an event history binomial model. In the binomial model, per each fly for each day, 0 coded for flies alive and 1 for dead. Protein and protein$^2$ are mean centered to standard deviation of 1. The model included random effects of Individual ID (posterior mean = 0.03 (95% credible interval (CI) = $7.56 \times 10^{-10}$ to 0.11), effective sample size = 1013) and Experimental day (posterior mean = 2.38 (95% CI = 1.61 to 3.25), effective sample size = 1000). Significant results below significance level $\alpha = 0.05$ are bolded.

| | Posterior mean | l-95% CI | u-95% CI | Effective sample size | pMCMC |
|---|---|---|---|---|---|
| **Intercept** | **-5.46** | **-5.89** | **-5.08** | **1000** | **<0.001** |
| Injury treatment | 0.14 | -0.32 | 0.57 | 1000 | 0.54 |
| **Infection treatment** | **0.66** | **0.28** | **1.10** | **1000** | **0.002** |
| Protein | 0.02 | -0.13 | 1.17 | 1000 | 0.82 |
| **Protein$^2$** | **0.48** | **0.26** | **0.71** | **1111** | **<0.001** |
| Injury:Protein | -0.08 | -0.30 | 0.15 | 1000 | 0.45 |
| **Infection:Protein** | **-0.31** | **-0.57** | **-0.10** | **1000** | **0.004** |
| Injury:Protein$^2$ | -0.16 | -0.51 | 0.18 | 1000 | 0.36 |
| Infection:Protein$^2$ | -0.01 | -0.33 | 0.30 | 1000 | 0.99 |

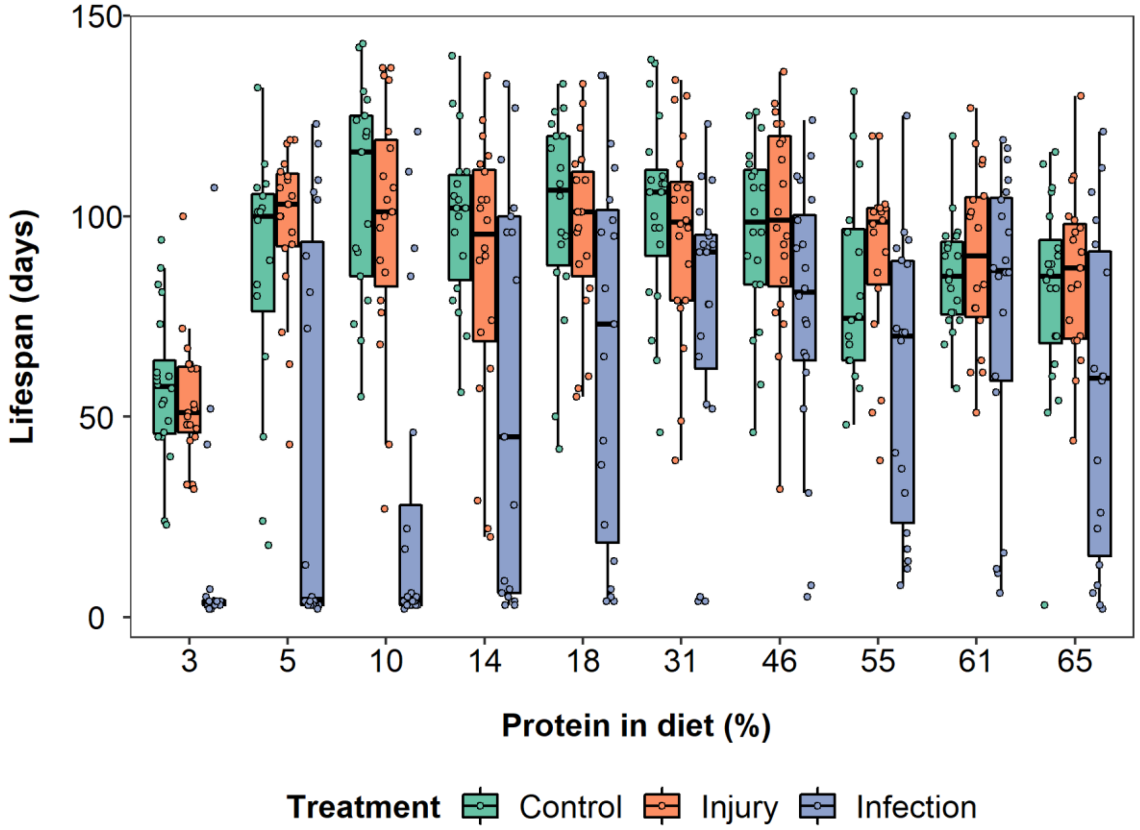


**Figure S3:** Effects of protein restriction on the lifespan of flies infected with a bacterial pathogen (blue bars and data points), injured by a pinprick (orange bars and data points) or with no treatment (green bars and data points). Data are observed lifespans (filled circles), where lines in the box plots indicate median lifespan (50% quantile), boxes are the interquartile range (25% to 75% quantiles) and whiskers are minimum or maximum quartiles (25% - 1.5 x interquartile range, 75% + 1.5 x interquartile range).

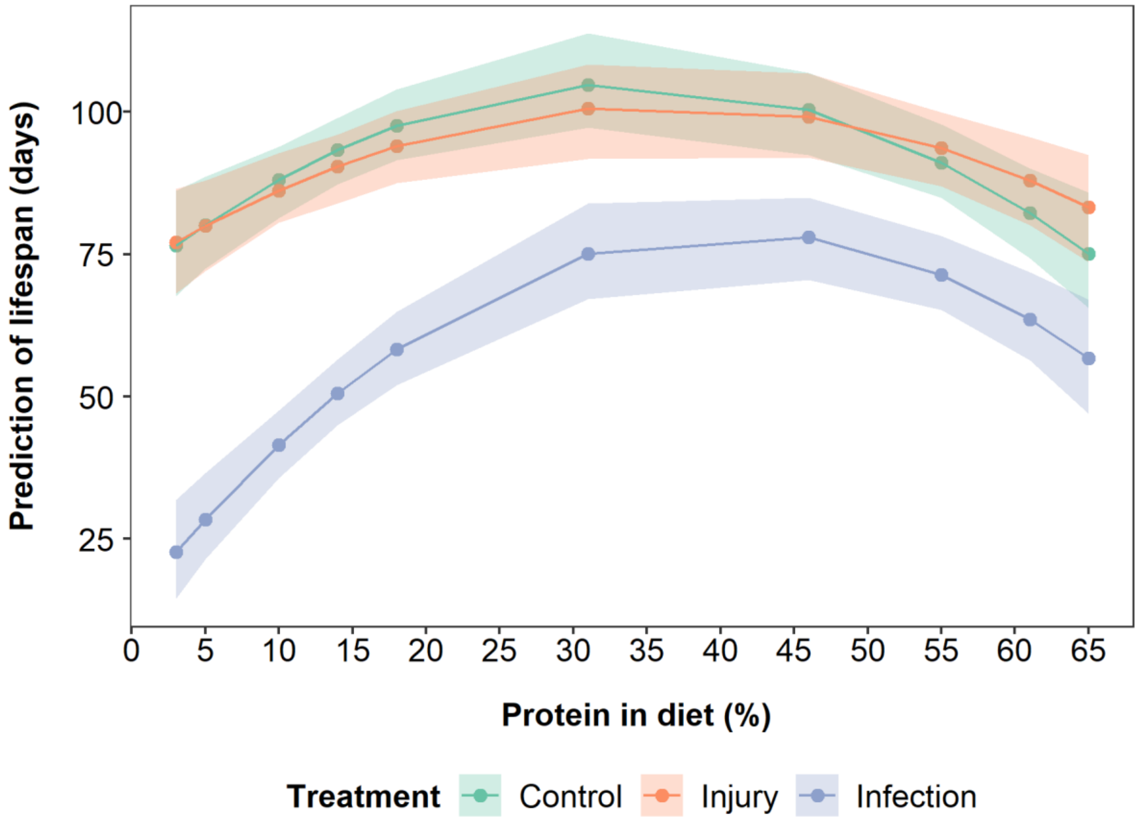


**Figure S4:** Model predictions of the effects of protein restriction on lifespan of flies infected with a bacterial pathogen (blue data points and lines), injured by a pinprick (orange data points and lines) or with no treatment (green data points and lines). Shaded areas are 95% 95% credible intervals. Protein and protein$^2$ are mean centered to standard deviation of 1.

**Table S3:** Model summary of effects of protein restriction and stress treatments on lifespan. Protein and protein$^2$ are mean centered to standard deviation of 1. The model included random effects of Individual ID (posterior mean = 0.028 (7.56 x $10^{-10}$-0.11), effective sample size = 1013) and Experimental day (posterior mean = 2.38 (1.61-3.25), effective sample size = 1000). Significant results below significance level α = 0.05 are bolded.

| | Posterior mean | l-95% CI | u-95% CI | Effective sample size | pMCMC |
|---|---|---|---|---|---|
| **Intercept** | **104.76** | **97.22** | **113.77** | **1000** | **<0.001** |
| Injury treatment | -4.17 | -15.32 | 7.55 | 1107 | 0.48 |
| **Infection treatment** | **-29.83** | **-41.32** | **-17.72** | **1000** | **<0.001** |
| Protein | 3.83 | -0.16 | -9.21 | 1000 | 0.09 |
| **Protein$^2$** | **-15.79** | **-22.55** | **-8.90** | **1108** | **<0.001** |
| Injury:Protein | 1.57 | -5.97 | 7.50 | 1000 | 0.65 |
| **Infection:Protein** | **14.31** | **7.66** | **20.99** | **1000** | **<0.001** |
| Injury:Protein$^2$ | 4.47 | -5.33 | 14.45 | 1158 | 0.39 |
| Infection:Protein$^2$ | -4.44 | -14.45 | 6.06 | 1000 | 0.40 |

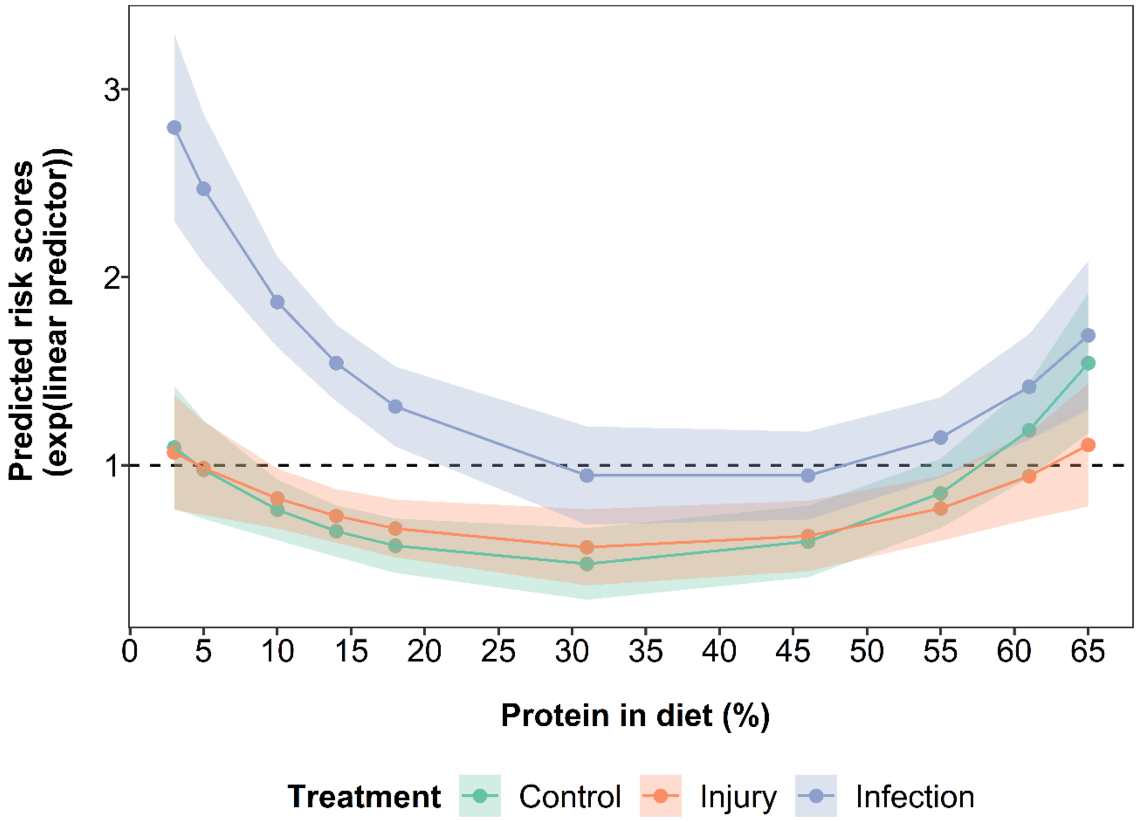


**Figure S5:** Model predictions for the effects of protein restriction on survival of flies infected with a bacterial pathogen (blue data points and lines), injured by a pinprick (orange data points and lines) or with no treatment (green data points and lines). $y = 1$ line shows no change in risk ratio, i.e. treatment would have no effect compared to baseline hazard. Shaded areas are 95% confidence intervals. Protein and protein$^2$ are mean centered to standard deviation of 1.

**Table S4:** Cox proportional hazard regression model summary of effects of protein restriction and stress treatments on survival (n = 600, number of deaths = 573, concordance = 0.662, $R^2$ = 0.142, Wald test = 97.98). Protein and protein$^2$ are mean centered to standard deviation of 1. Significant results below significance level $\alpha = 0.05$ are bolded.

| | coef | exp(coef) | se(coef) | z | Pr (>\|z\|) |
|---|---|---|---|---|---|
| Injury treatment | 0.17 | 1.19 | 0.20 | 0.84 | 0.40 |
| **Infection treatment** | **0.69** | **1.99** | **0.21** | **3.36** | **<0.001** |
| Protein | -0.03 | 0.97 | 0.09 | -0.29 | 0.77 |
| **Protein$^2$** | **0.54** | **1.72** | **0.12** | **4.28** | **<0.001** |
| Injury:Protein | -0.06 | 0.94 | 0.12 | -0.52 | 0.60 |
| **Infection:Protein** | **-0.29** | **0.75** | **0.12** | **-2.43** | **0.01** |
| Injury:Protein$^2$ | -0.18 | 0.83 | 0.18 | -1.03 | 0.30 |
| Infection:Protein$^2$ | -0.07 | 0.93 | 0.18 | -0.41 | 0.68 |

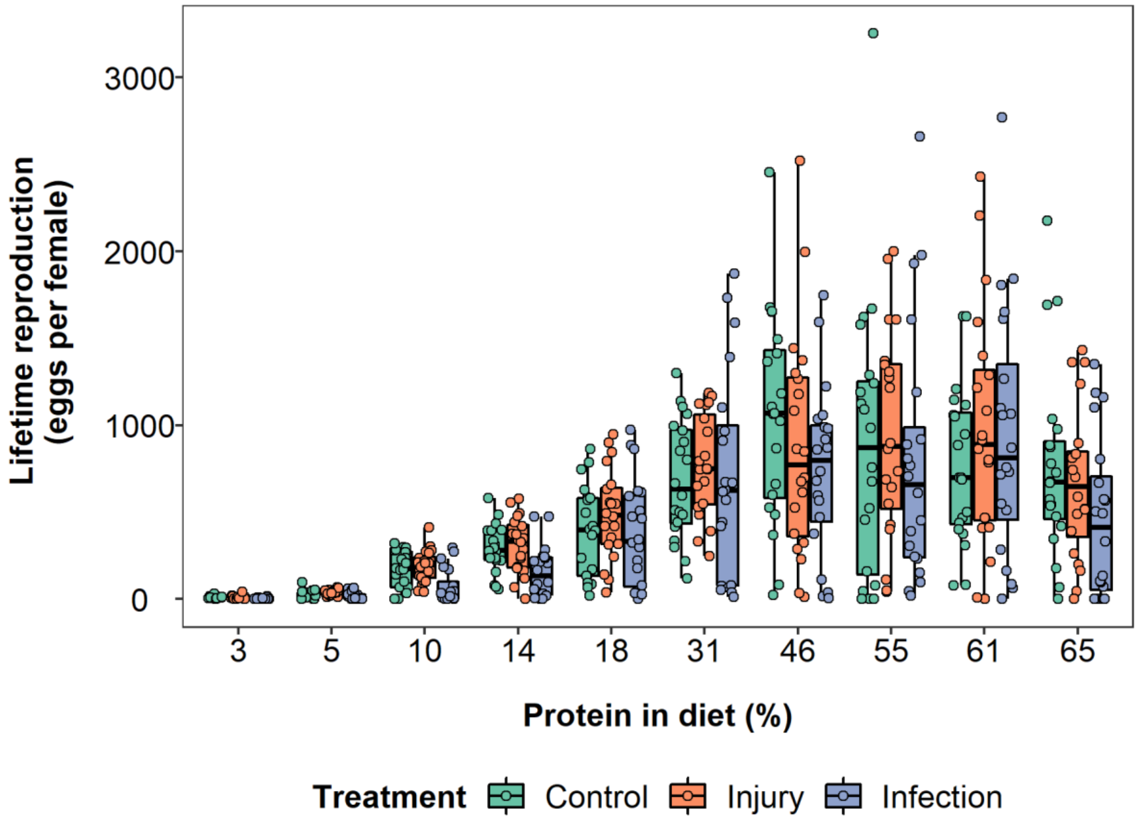


**Figure S6:** Effect of protein restriction on the lifetime egg production of flies infected with a bacterial pathogen (blue lines and data points), injured by a pinprick (orange lines and data points) or with no treatment (green lines and data points). The lines in the box plots indicate median lifespan (50% quantile), the boxes are the interquartile range (25% to 75% quantiles) and the whiskers are minimum or maximum quartiles (25% - 1.5 x interquartile range, 75% + 1.5 x interquartile range).

**Table S5:** Model summary of effects of protein restriction and stress treatments on lifetime eggs produced. Protein and protein$^2$ are mean centered to standard deviation of 1. Significant results below significance level $\alpha = 0.05$ are bolded.

| | Posterior mean | l-95% CI | u-95% CI | Effective sample size | pMCMC |
|---|---|---|---|---|---|
| **Intercept** | **6.55** | **6.17** | **6.92** | **1205** | **<0.001** |
| Injury treatment | 0.19 | -0.34 | 0.72 | 1000 | 0.49 |
| Infection treatment | -0.33 | -0.90 | 0.16 | 1330 | 0.264 |
| **Protein** | **1.45** | **1.23** | **1.64** | **1000** | **<0.001** |
| **Protein$^2$** | **-1.36** | **-1.68** | **-1.02** | **1000** | **<0.001** |
| Injury:Protein | -0.09 | -0.39 | -1.02 | 1000 | 0.60 |
| **Infection:Protein** | **0.47** | **0.16** | **0.77** | **1000** | **0.01** |
| Injury:Protein$^2$ | -0.02 | -0.45 | 0.46 | 1000 | 0.93 |
| **Infection:Protein$^2$** | **-0.47** | **-0.93** | **-0.04** | **1000** | **0.04** |

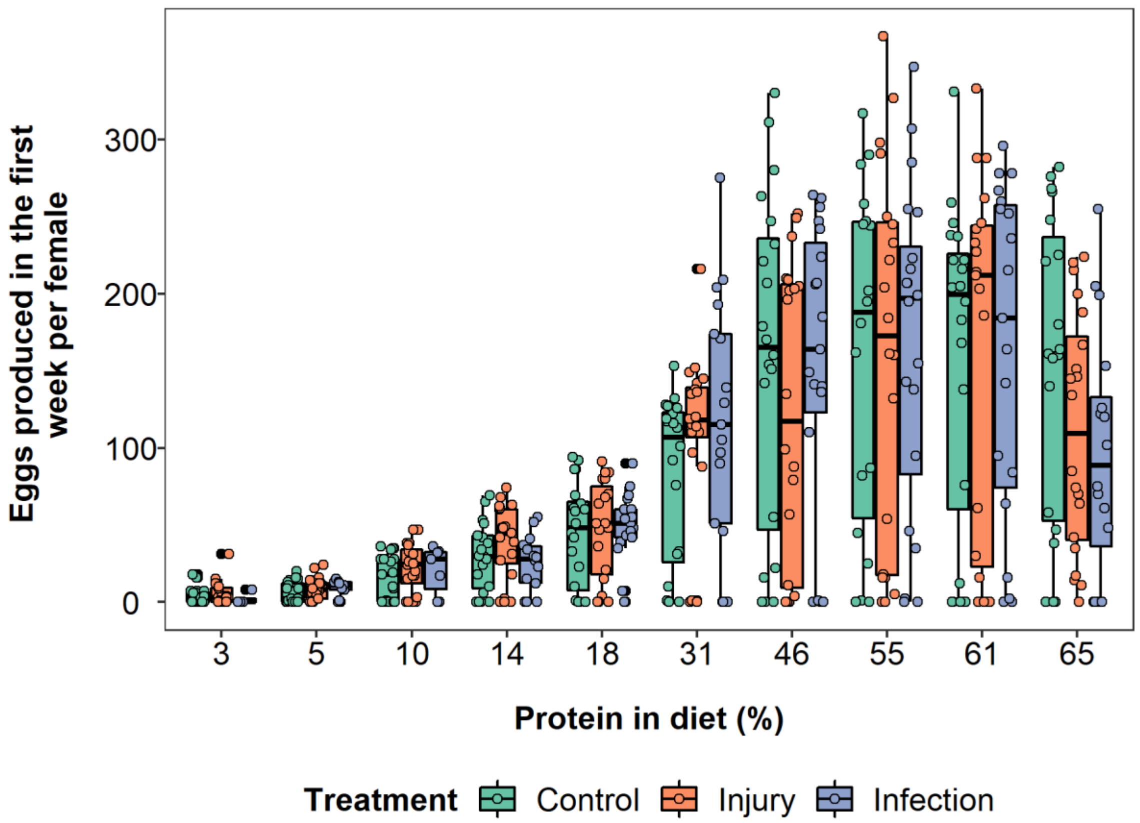


**Figure S7:** Effect of protein restriction on the early-life egg production of flies infected with a bacterial pathogen (blue lines and data points), injured by a pinprick (orange lines and data points) or with no treatment (green lines and data points). Early-egg production consists of the first seven days of egg production without the first day (see methods). The lines in the box plots indicate median number of eggs produced (50% quantile), boxes are the interquartile range (25% to 75% quantiles) and whiskers are minimum or maximum quartiles (25% - 1.5 x interquartile range, 75% + 1.5 x interquartile range).

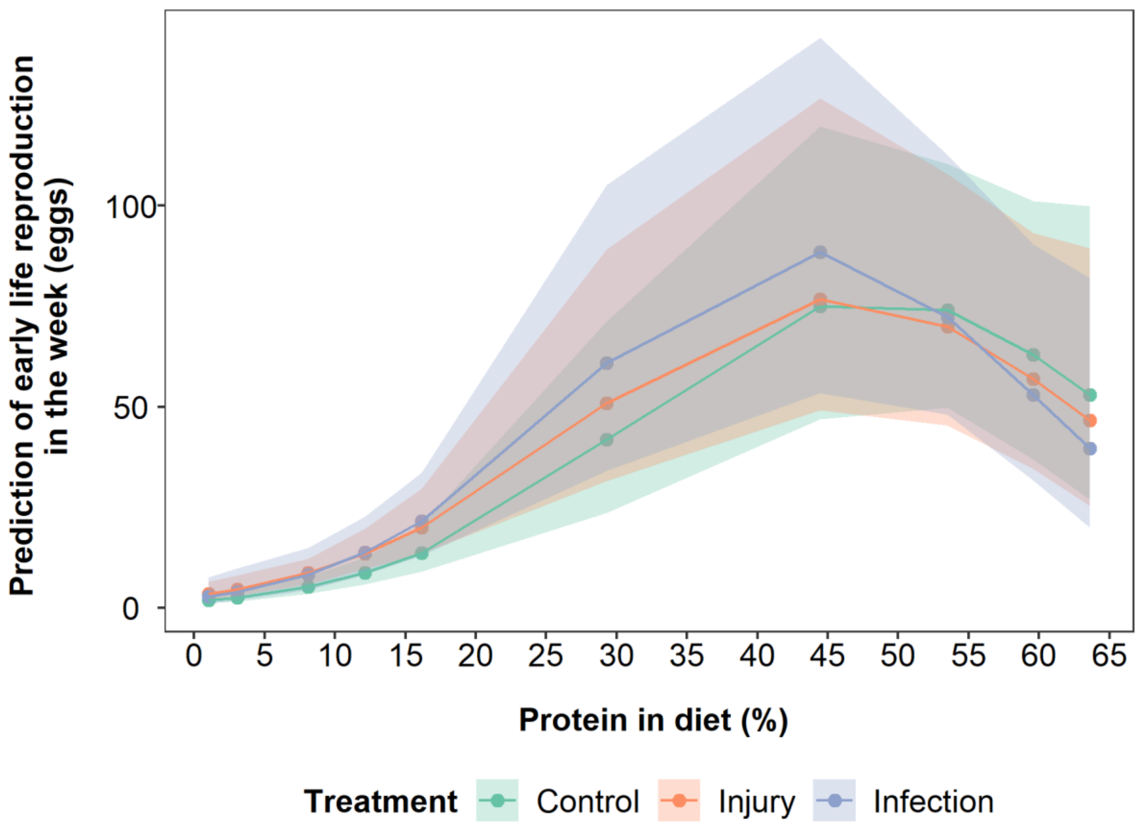


**Figure S8:** Model predictions of the effect of protein restriction on the early-life egg production of flies infected with a bacterial pathogen (blue data points and lines), injured by a pinprick (orange data points and lines) or with no treatment (green data points and lines). Early-life egg production consists of the first seven days of egg production without the first day (see methods). Shaded areas are 95% 95% credible intervals. Protein and protein$^2$ are mean centered to standard deviation of 1.

**Table S6:** Model summary of effect of protein restriction and stress treatment on early-life egg production (first week discounting the first day, see methods). Protein and protein$^2$ are mean centered to standard deviation of 1. Significant results below significance level $\alpha = 0.05$ are bolded.

| | Posterior mean | l-95% CI | u-95% CI | Effective sample size | pMCMC |
|---|---|---|---|---|---|
| **Intercept** | **3.82** | **3.30** | **4.41** | **1156** | **<0.001** |
| Injury treatment | 0.18 | -0.56 | 0.92 | 1000 | 0.69 |
| Infection treatment | 0.36 | -0.37 | 1.16 | 1330 | 0.37 |
| **Protein** | **1.34** | **1.06** | **1.64** | **1000** | **<0.001** |
| **Protein$^2$** | **-0.86** | **-1.34** | **-0.41** | **1149** | **<0.001** |
| Injury:Protein | -0.29 | -0.69 | 0.10 | 1000 | 0.17 |
| Infection:Protein | -0.24 | -0.74 | 0.20 | 1000 | 0.32 |
| Injury:Protein$^2$ | 0.05 | -0.60 | 0.66 | 1000 | 0.86 |
| Infection:Protein$^2$ | -0.14 | -0.85 | 0.57 | 1198 | 0.74 |

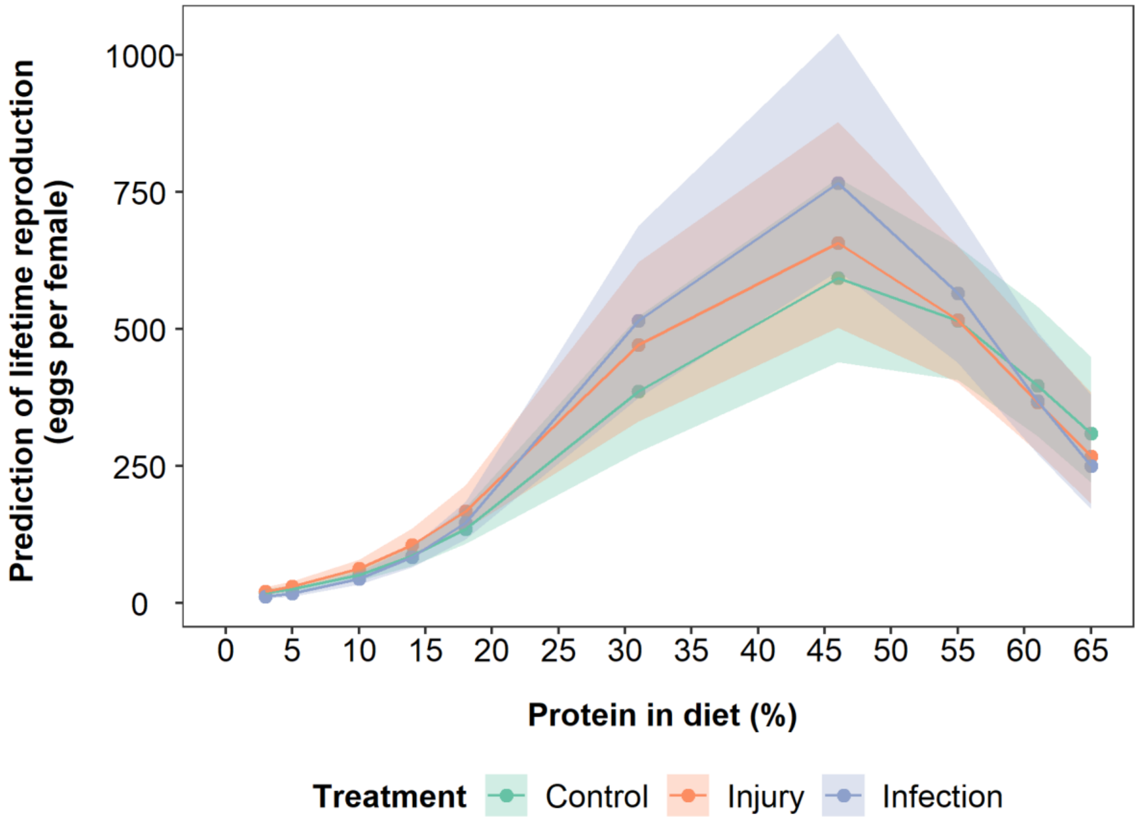


**Figure S9:** Model predictions of the effects of protein restriction on the lifetime number of eggs produced by flies infected with a bacterial pathogen (blue data points and lines), injured by a pinprick (orange data points and lines) or with no treatment (green data points and lines), when accounting for lifespan (mean centred). Shaded areas are 95% credible intervals. Protein and protein$^2$ are mean centered to standard deviation of 1.

**Table S7:** Model summary of effects of protein restriction and stress treatments on lifetime eggs produced. Mean centered lifespan is added as a fixed effect to remove the effect of lifespan on reproduction. Protein and protein$^2$ are mean centered to standard deviation of 1. Significant results below significance level α = 0.05 are bolded.

| | Posterior mean | l-95% CI | u-95% CI | Effective sample size | pMCMC |
|---|---|---|---|---|---|
| **Intercept** | **5.94** | **5.64** | **6.28** | **1000** | **<0.001** |
| Injury treatment | 0.20 | -0.19 | 0.67 | 1000 | 0.36 |
| Infection treatment | 0.29 | -0.18 | 0.72 | 1000 | 0.21 |
| **Protein** | **1.31** | **1.15** | **1.50** | **1119** | **<0.001** |
| **Protein$^2$** | **-0.97** | **-1.24** | **-0.70** | **1000** | **<0.001** |
| **Lifespan** | **0.93** | **0.83** | **1.04** | **1000** | **<0.001** |
| Injury:Protein | -0.81 | -0.33 | 0.17 | 1000 | 0.52 |
| Infection:Protein | 0.18 | -0.07 | 0.45 | 1000 | 0.17 |
| Injury:Protein$^2$ | -0.10 | -0.45 | 0.28 | 1000 | 0.60 |
| Infection:Protein$^2$ | -0.34 | -0.71 | 0.05 | 1000 | 0.08 |

**AGEING:**

DAILY EGG PRODUCTION:

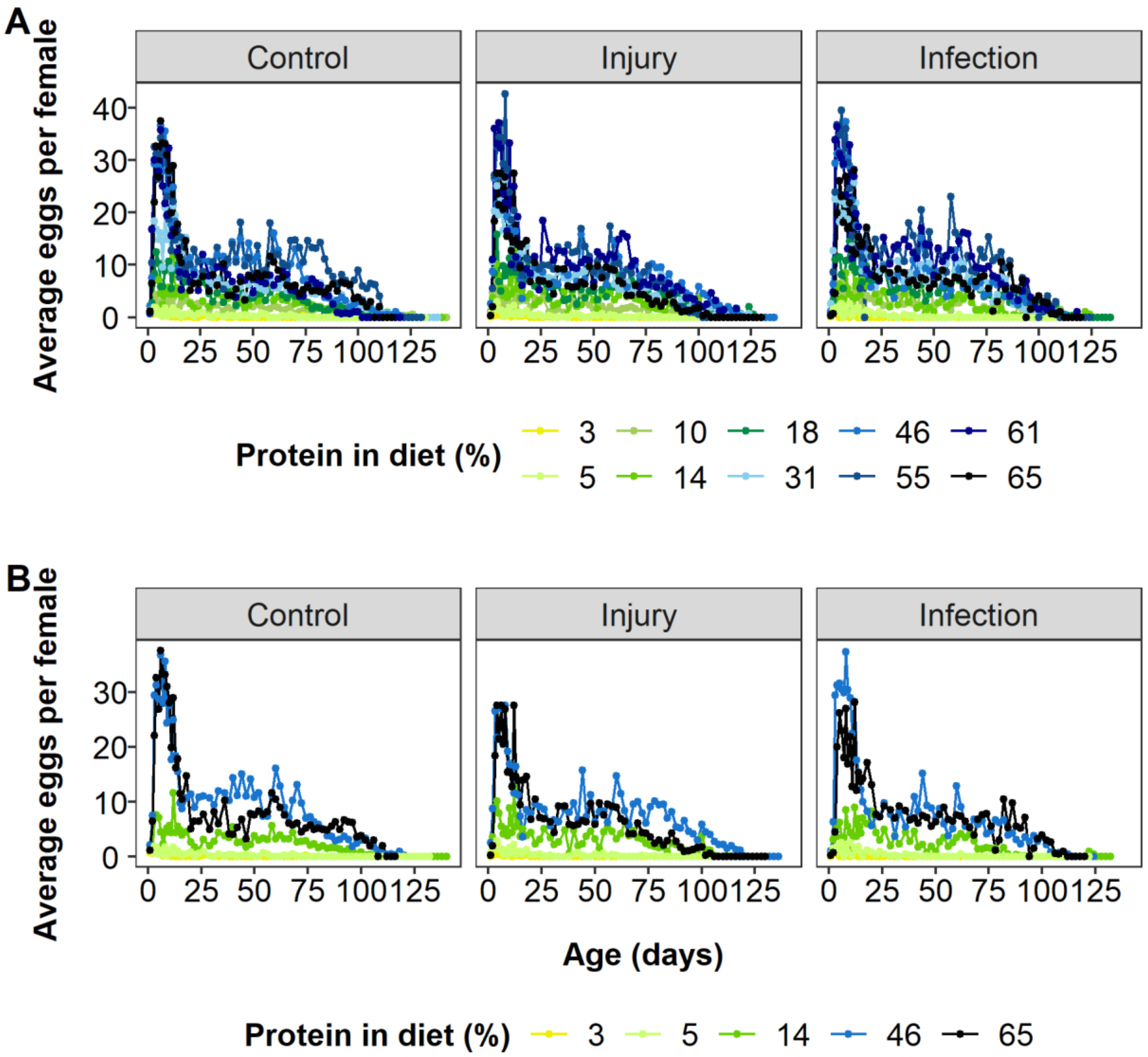


**Figure S10:** The pattern of ageing in egg production for each protein restriction diet for flies infected with a bacterial pathogen ("Infection"), injured by a pinprick ("Injury") or with no treatment ("Control"). The average eggs laid per day across all flies per diet and stress treatment is plotted and the associated errors have been removed for clarity. (A) All diets for each stress treatment; (B) A subset of protein restriction diets to illustrate the effects of protein restriction with low (yellow line) intermediate (blue line) and high protein content (black line).

**Table S8:** Model summary of effects of protein restriction, age and stress treatment for daily egg production on flies. Protein, protein$^2$, age, age$^2$ and lifespan are mean centered to standard deviation of 1. The model included random effects of Individual ID (posterior mean = 1.76 (95% CI = 1.53 to 2.01), effective sample size = 1334). Lifespan (mean centered) is included to account for selective disappearance. Significant results below significance level $\alpha = 0.05$ are bolded.

| | Posterior mean | l-95% CI | u-95% CI | Effective sample size | pMCMC |
|---|---|---|---|---|---|
| **Intercept** | **1.61** | **1.23** | **1.98** | **1334** | **<0.001** |
| Injury treatment | 0.19 | -0.35 | 0.68 | 1193 | 0.48 |
| Infection treatment | -0.11 | -0.66 | 0.47 | 1334 | 0.72 |
| **Protein** | **1.31** | **1.12** | **1.52** | **1334** | **<0.001** |
| **Protein$^2$** | **-1.51** | **-1.81** | **-1.19** | **1334** | **<0.001** |
| **Age** | **-0.32** | **-0.40** | **-0.23** | **1222** | **<0.001** |
| **Age$^2$** | **-0.52** | **-0.59** | **-0.44** | **1334** | **<0.001** |
| **Lifespan** | **0.21** | **0.11** | **0.31** | **1334** | **<0.001** |
| Injury:Protein | 0.08 | -0.22 | 0.35 | 1334 | 0.58 |
| Infection:Protein | -0.21 | -0.54 | 0.11 | 1334 | 0.22 |
| Injury:Protein$^2$ | -0.05 | -0.51 | 0.36 | 1477 | 0.82 |
| **Infection:Protein$^2$** | **051** | **0.04** | **0.97** | **1334** | **0.03** |
| **Injury:Age** | **0.13** | **0.01** | **0.24** | **1334** | **0.03** |
| **Infection:Age** | **-0.29** | **-0.42** | **-0.14** | **1334** | **<0.001** |
| Injury:Age$^2$ | 0.08 | -0.03 | 0.19 | 1063 | 0.15 |
| **Infection:Age$^2$** | **0.27** | **0.13** | **0.41** | **1334** | **0.002** |
| Protein:Age | 0.02 | -0.04 | 0.08 | 1334 | 0.56 |
| Protein:Age$^2$ | -0.04 | -0.09 | 0.02 | 1660 | 0.22 |
| **Protein$^2$:Age** | **-0.24** | **-0.32** | **-0.16** | **1334** | **<0.001** |
| **Protein$^2$:Age$^2$** | **0.30** | **0.22** | **0.38** | **1116** | **<0.001** |
| **Injury:Protein:Age** | **0.14** | **0.05** | **0.22** | **1334** | **0.002** |
| **Infection:Protein:Age** | **0.11** | **0.001** | **0.20** | **1334** | **0.04** |
| **Injury:Protein:Age$^2$** | **-0.14** | **-0.22** | **-0.06** | **1334** | **<0.001** |

| | | | | | |
|---|---|---|---|---|---|
| Infection:Protein:Age$^2$ | 0.02 | -0.09 | 0.13 | 1334 | 0.64 |
| **Injury:Protein$^2$:Age** | **-0.12** | **-0.23** | **-0.005** | **1360** | **0.04** |
| **Infection:Protein$^2$:Age** | **0.22** | **0.09** | **0.35** | **1334** | **0.005** |
| **Injury:Protein$^2$:Age$^2$** | **-0.13** | **-0.25** | **-0.02** | **1175** | **0.03** |
| **Infection:Protein$^2$:Age$^2$** | **-0.37** | **-0.51** | **-0.23** | **1334** | **<0.001** |

SMURF ASSAY:

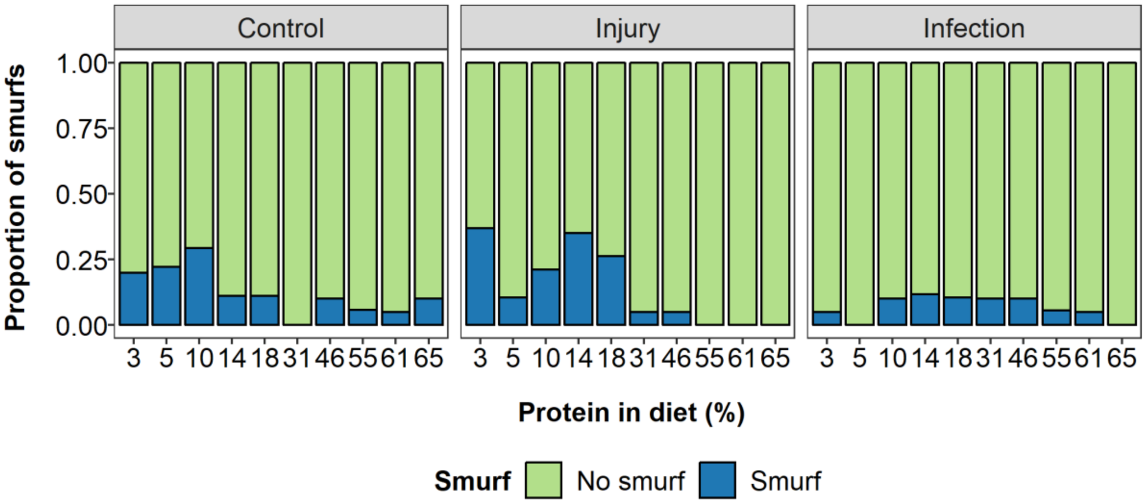


**Figure S11:** Effects of protein restriction on proportion of smurfs (blue bars) or no smurfs (green bars) across life of flies infected with a bacterial pathogen (“Infection”, N = 23), injured by a pinprick (“Injury”, N = 25) or with no treatment (“Control”, N = 15).

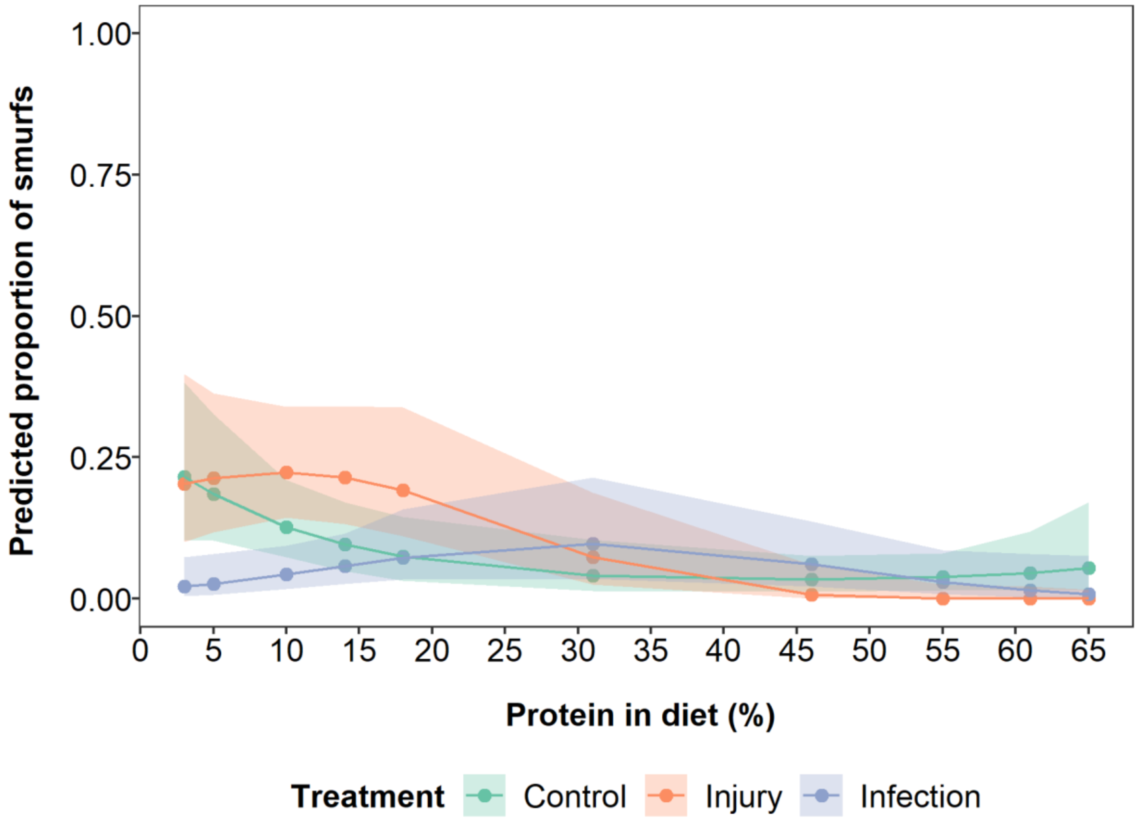


**Figure S12:** Model predictions of the effect of protein restriction on the proportion of flies developing into a smurf of flies infected with a bacterial pathogen (blue data points and lines), injured by pinprick (orange data points and lines) or with no treatment (green data points and lines). Protein and protein$^2$ are mean centered to standard deviation of 1. Shaded areas are 95% credible intervals.

**Table S9:** Model summary of effects of protein restriction and stress treatment on proportion of flies developing into a smurf. Protein and protein$^2$ are mean centered to standard deviation of 1. Significant results below significance level $\alpha = 0.05$ are bolded.

| | Posterior mean | l-95% CI | u-95% CI | Effective sample size | pMCMC |
|---|---|---|---|---|---|
| **Intercept** | **-3.13** | **-4.21** | **-2.07** | **1000** | **<0.001** |
| Injury treatment | 0.63 | -0.81 | 2.27 | 1000 | 0.37 |
| Infection treatment | 0.91 | -0.59 | 2.30 | 1000 | 0.23 |
| **Protein** | **-0.75** | **-1.24** | **-0.21** | **1000** | **0.004** |
| Protein$^2$ | 0.63 | -0.26 | 1.41 | 1000 | 0.15 |
| **Injury:Protein** | **-1.96** | **-4.07** | **-0.11** | **1000** | **0.01** |
| Infection:Protein | 0.73 | -0.20 | 1.71 | 1000 | 0.10 |
| **Injury:Protein$^2$** | **-2.09** | **-4.22** | **-0.47** | **1000** | **0.14** |
| **Infection:Protein$^2$** | **-1.73** | **-3.13** | **-0.37** | **1000** | **0.01** |

NEGATIVE GEOTAXIS (NG):

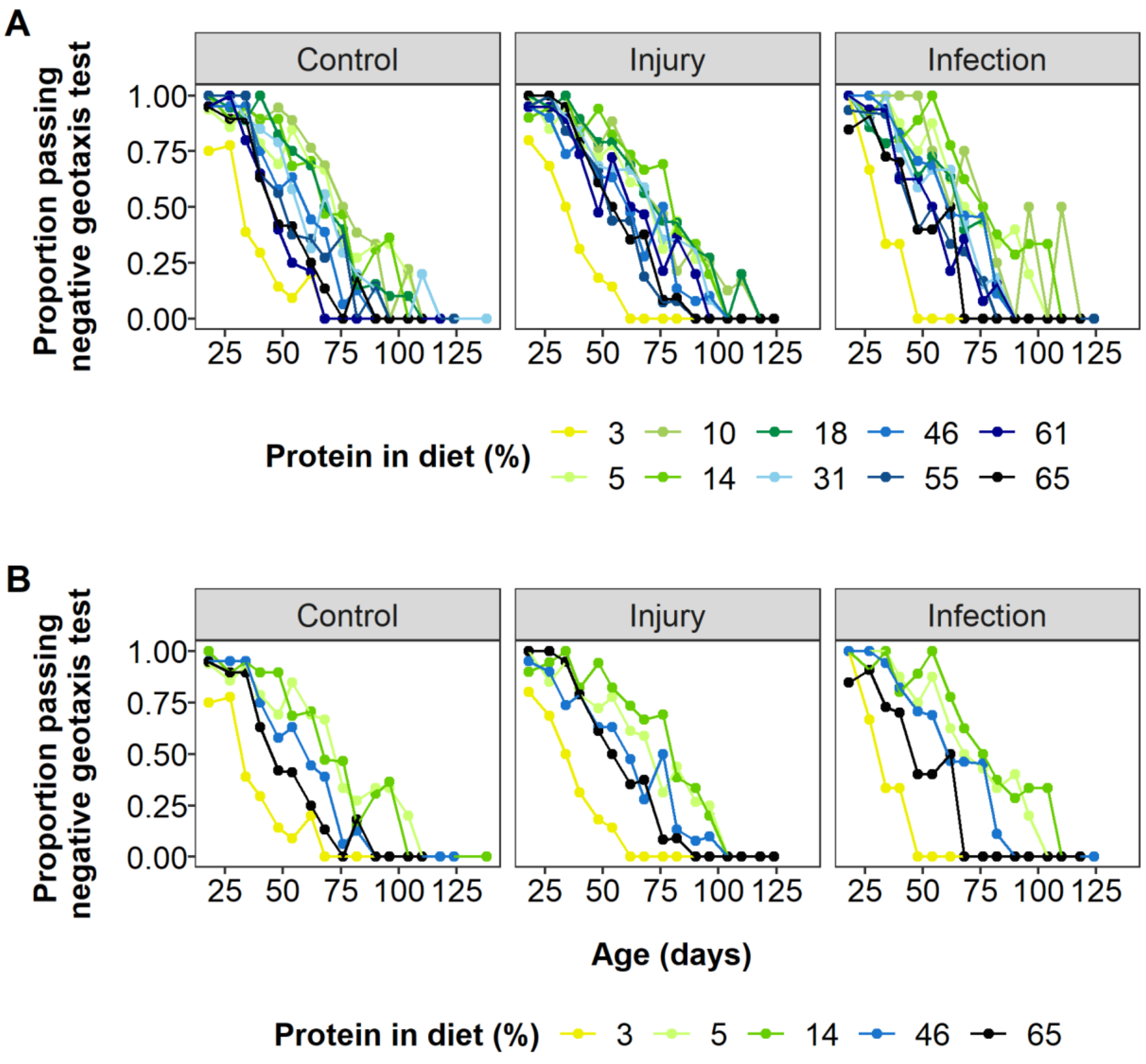


**Figure S13:** Effects of protein restriction on the proportion of flies passing the negative geotaxis test under 60 seconds per week of flies infected with a bacterial pathogen ("Infection"), injured by a pinprick ("Injury"), or with no treatment ("Control") (A). For ease of interpretation, a subset of diets is shown in (B) to illustrate the effects of protein restriction with low (yellow line) intermediate (pale blue line) and high protein content (black line).

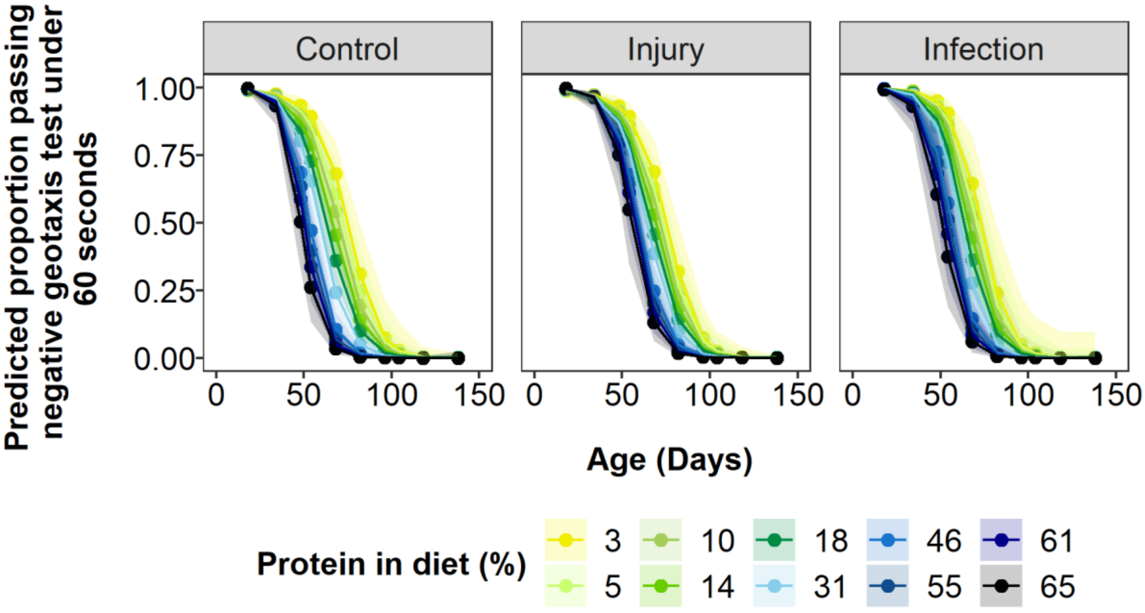


**Figure S14:** Model predictions of the effect of protein restriction and age on proportion passing negative geotaxis test under 60 seconds per week with flies infected with a bacterial pathogen ("Infection"), injured by pinprick ("Injury") or with no treatment ("Control"). Shaded areas are 95% credible intervals. Protein, protein$^2$ and lifespan are mean centered to standard deviation of 1.

**Table S10:** Model summary of effects of protein restriction, age and stress treatment for passing negative geotaxis test under 60 seconds. Protein, protein$^2$, age, age$^2$ and lifespan are mean centered to standard deviation of 1. Lifespan (mean centered) is included to account for selective disappearance. The model included the random effect of Individual ID (posterior mean = 3.03 (95% CI = 2.35 to 3.73), effective sample size = 1000). Significant results below significance level α = 0.05 are bolded.

| | Posterior mean | l-95% CI | u-95% CI | Effective sample size | pMCMC |
|---|---|---|---|---|---|
| **Intercept** | **1.01** | **0.39** | **1.63** | **892.4** | **0.002** |
| Injury treatment | 0.52 | -0.38 | 1.37 | 1000 | 0.23 |
| Infection treatment | 0.35 | -0.59 | 1.32 | 1000 | 0.49 |
| **Protein** | **-0.65** | **-1.01** | **-0.32** | **1000** | **<0.001** |
| **Protein$^2$** | **-0.70** | **-1.21** | **-0.21** | **1060** | **0.01** |
| **Age** | **-3.57** | **-4.04** | **-3.07** | **1000** | **<0.001** |
| Age$^2$ | -0.13 | -0.61 | 0.36 | 902.6 | 0.58 |
| **Lifespan** | **0.84** | **0.64** | **1.02** | **1000** | **<0.001** |
| Injury:Protein | 0.38 | -0.07 | 0.86 | 1197.5 | 0.12 |
| Infection:Protein | 0.20 | -0.40 | 0.81 | 1000 | 0.53 |
| Injury:Protein$^2$ | 0.10 | -0.60 | 0.78 | 1102.8 | 0.78 |
| Infection:Protein$^2$ | -0.04 | -0.76 | 0.89 | 1101.4 | 0.93 |
| Injury:Age | 0.48 | -0.15 | 1.17 | 1098.6 | 0.17 |
| Infection:Age | -0.17 | -1.03 | 0.51 | 1039.8 | 0.69 |
| Injury:Age$^2$ | -0.30 | -0.86 | 0.43 | 1000 | 0.37 |
| Infection:Age$^2$ | -0.26 | -1.12 | 0.47 | 1000 | 0.51 |
| **Protein:Age** | **-0.78** | **-1.06** | **-0.49** | **1108.6** | **<0.001** |
| Protein:Age$^2$ | 0.16 | -0.13 | 0.42 | 1000 | 0.27 |
| Protein$^2$:Age | 0.06 | -0.33 | 0.48 | 1124.5 | 0.79 |
| Protein$^2$:Age$^2$ | 0.15 | -0.28 | 0.50 | 1000 | 0.45 |
| Injury:Protein:Age | 0.28 | -0.11 | 0.63 | 1000 | 0.16 |
| Infection:Protein:Age | 0.37 | -0.10 | 0.91 | 1132.2 | 0.17 |
| Injury:Protein:Age$^2$ | -0.17 | -0.54 | 0.19 | 1045.3 | 0.40 |

| | | | | | |
|---|---|---|---|---|---|
| Infection:Protein:Age$^2$ | -0.35 | -0.83 | 0.10 | 1227 | 0.14 |
| Injury:Protein$^2$:Age | -0.17 | -0.72 | 0.39 | 1000 | 0.54 |
| Infection:Protein$^2$:Age | -0.03 | -0.63 | 0.75 | 1000 | 0.95 |
| Injury:Protein$^2$:Age$^2$ | 0.09 | -0.45 | 0.60 | 1000 | 0.73 |
| Infection:Protein$^2$:Age$^2$ | 0.13 | -0.56 | 0.80 | 1000 | 0.69 |
